# Dynamic ParB-DNA interactions initiate and maintain a partition condensate for bacterial chromosome segregation

**DOI:** 10.1101/2023.06.25.546419

**Authors:** Miloš Tišma, Richard Janissen, Hammam Antar, Alejandro Martin Gonzalez, Roman Barth, Twan Beekman, Jaco van der Torre, Davide Michieletto, Stephan Gruber, Cees Dekker

## Abstract

In most bacteria, chromosome segregation is driven by the ParAB*S* system where the CTPase protein ParB loads at the *parS* site to trigger the formation of a large partition complex. Here, we present *in vitro* studies of the partition complex for *Bacillus subtilis* ParB, using single-molecule fluorescence microscopy and AFM imaging to show that transient ParB-ParB bridges are essential for forming DNA condensates. Molecular Dynamics simulations confirm that condensation occurs abruptly at a critical concentration of ParB and show that multimerization is a prerequisite for forming the partition complex. Magnetic tweezer force spectroscopy on mutant ParB proteins demonstrates that CTP hydrolysis at the N-terminal domain is essential for DNA condensation. Finally, we show that transcribing RNA polymerases can steadily traverse the ParB-DNA partition complex. These findings uncover how ParB forms a stable yet dynamic partition complex for chromosome segregation that induces DNA condensation and segregation while enabling replication and transcription.

## Introduction

Precise chromosome segregation at each cell cycle is important for the stable propagation of all life forms. In most bacteria, the ParAB*S* system is the main component ensuring such faithful segregation of chromosomes^1^. This system consists of an ATP-hydrolase partition protein A (ParA), a CTP-hydrolase partition protein B (ParB), and a 16-bp centromeric sequence *parS* that is typically located near the replication origin on the genome ^2–5^. ParB proteins can load at the *parS* sequence and subsequently assemble into a higher-order nucleoprotein complex that is referred to as ‘the partition complex’ ^6–9^. Following replication, the two partition complexes interact with a ParA gradient along the cylindrical cell axis to segregate the nascent origins of bacterial chromosomes ^10, 11^. Bacterial SMC proteins are recruited to the partition complex ^12–14^ to further increase the fidelity of the segregation process ^14–18^. The formation of a functional partition complex is essential to ensure a correct distribution of chromosomes during cell division.

Several models have been proposed for the molecular structure of the partition complex (reviewed in detail by Jalal and Le ^19^), including ‘bridging and condensing’ ^8, 20, 21^, ‘nucleation and caging’ ^22, 23^, and recent work indicating liquid-liquid phase-separated droplets ^24, 25^. In essence, all these models rely on the initial nucleation of ParB proteins at the *parS* site followed by some ‘self-self’ interactions by ParB proteins to bring distal DNA elements together to form a partition complex. Many older models that relied on *in vitro* insights required revision after it was shown that ParB proteins utilize CTP to load at a *parS*-site and able to spread over multiple kilobases ^3, 4, 26^. ParB proteins were shown to condense DNA in the presence of CTP, and a new model combining one-dimensional sliding and ParB-ParB bridging has been proposed ^27^, in which ParB dimers form a clamp that binds two CTP molecules upon loading at the *parS* site, whereupon they lose the affinity to *parS* and slide along the DNA to spread to adjacent DNA regions ^4, 28, 29^. After some time, the ParB clamp is hypothesized to open either at the C- or N-terminus, which enables to bring two distant DNA segments into proximity through *in-trans* bridging interactions ^27^. This model integrates the DNA ‘clamping and sliding’ hypothesis in the presence of CTP with previous models that considered ‘caging’ or ‘bridging’ of ParB proteins ^8, 9, 20, 21, 30^.

Many questions remain, however, on the formation and maintenance of the partition complex. For example, what is the structure and dynamics of the partition complex? How is the condensation of the complex initiated? How does CTP hydrolysis regulate both the DNA binding and clamping, and the self-self interactions needed for bridge formation? Which protein domains are involved in bridge formation? While the necessity of both C- and N-terminal domains is clear ^20, 31^, it remains to be determined which conformational changes need to occur prior to bridge formation and DNA condensation ^4, 28, 29^. Moreover, partition complex formation via DNA condensation has never been reconstituted *in vitro* using a single *parS* site, while a single *parS* site was found to be sufficient for *in vivo* partition complex function^18, 32^.

To address these questions, we investigated the mechanism of ParB-DNA complex formation *in vitro* at the single-molecule level using purified *Bacillus subtilis* ParB proteins in Atomic Force Microscopy (AFM), Magnetic Tweezers (MT), and visualization by TIRF microscopy in a DNA-stretching assay ^33^. Furthermore, we performed Molecular Dynamics (MD) simulations using a coarse-grained DNA polymer and ParB models. Finally, we used *in vitro* transcription to clarify the debate if a ParB:DNA cluster affects transcription, and we observed that, remarkably, transcription by bacterial DNA-dependent RNA polymerase (RNAP) is not affected by ParB or condensed ParB:DNA clusters.

## Results

### ParB proteins condense DNA with a single *parS* site

The recent discovery that ParB proteins require CTP nucleotide to specifically load onto a DNA with a *parS* site (DNA*_parS_*) and perform their cellular function challenged many established models of ParB function ^3, 4, 34^. Most previous condensation studies used very high protein concentrations in order to show the occurrence of ParB condensation or DNA binding ^2, 8, 20, 21, 31^, and often no differences were found in the presence or absence of a *parS* site on the DNA substrate ^8, 20, 21^. Recent work by Balaguer *et al.* ^27^, however, showed that in the presence of CTP nucleotide, ParB proteins could induce condensation of a ∼6 kbp DNA at nanomolar concentrations, in line with the efficient loading of ParB. This study reported that at least 7 *parS* sites were necessary on the DNA for condensation to occur ^27^, which was unexpected given that *B. subtilis* mutants carrying a single *parS* site showed no defective phenotype in growth or ParB loading in vivo ^18, 32^.

To quantify the ability of ParB to drive condensation, we first used MT to study the interaction of *B. subtilis* ParB proteins with a long DNA substrate (∼14 kbp) that contained only a single *parS* site in the middle of the construct. Chemical groups attached to opposite ends of this DNA*_parS_* were used to tether one end of the DNA to a streptavidin-coated magnetic bead (through a biotin group) and the other end to an anti-DIG antibody-coated surface (through a digoxigenin group) (Fig. 1A, S1A, see Methods). We incrementally lowered the force that was applied to the magnetic bead from 7 to 0.05 pN (Fig. 1B – blue line) and recorded the change in end-to-end length of the attached DNA molecule.

**Figure 1.**
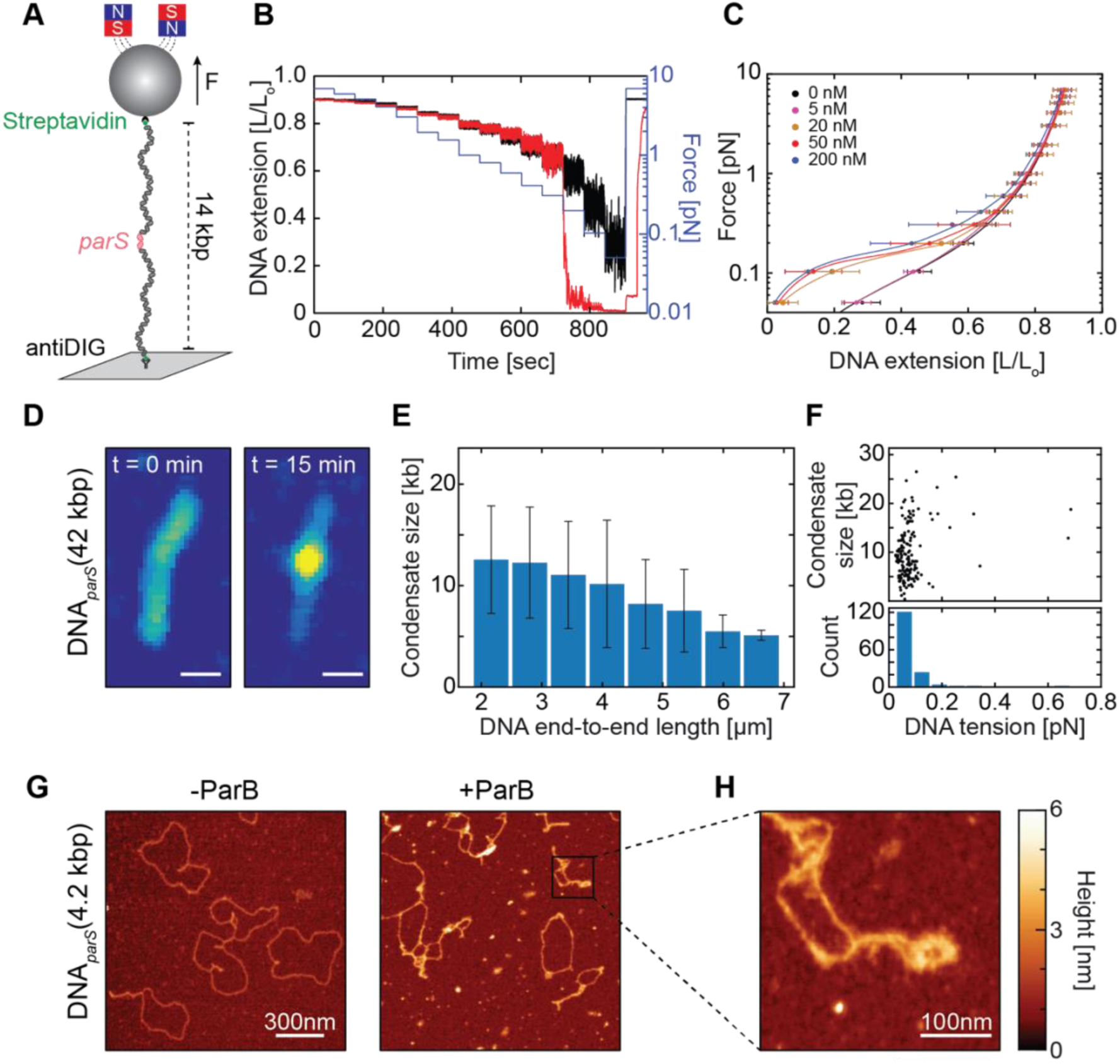
ParB proteins efficiently condense DNA in the presence of a single *parS* site. **(A)** Schematic representation of experimental MT assay, using a single *parS* site at the DNA*_parS_* molecule (see Methods). (**B)** DNA extension in the absence (black) and the presence (red) of 50 nM ParB protein at different forces applied to the magnetic bead (blue) over time. **C)** Average DNA*_parS_* extension (mean±SD) when lowering force (N = 35 for each ParB concentration) in the presence of different ParB concentrations. Black line represents a WLC fit to the 0 nM ParB control data. (**D)** Snapshot fluorescence images of a DNA*_parS_* molecule before and after 15 min incubation with 25 nM ParB. (**E)** The effect of DNA tethering length on final condensate size (mean±SD; N = 153), represented as kilobase pairs of the DNA that it contains. r = - 0.64 (Pearson correlation coefficient). (**F)** Tension that the formed ParB clusters of different sizes exert on the DNA fragments outside the cluster. N = 153. **G)** DNA condensation in the absence and presence of 7 nM ParB molecules. (**H)** Magnified image represents a high-resolution image of the ParB-DNA cluster. See also Figure S1.

Strikingly, in the presence of ParB and CTP, a single *parS* site on the DNA was found to be sufficient for DNA condensation, as evidenced by the abrupt reduction of the DNA end-to-end length (i.e., a rapid lowering of the bead height) when lowering the force below ∼0.3 pN (Fig. 1B - red line). This was not observed when ParB protein was not present (Fig. 1B - black line). Similarly, we did not observe any DNA condensation in the absence of CTP or in the absence of a *parS* site (Fig. S1B-C). We quantified the force-extension curves for different concentrations of ParB and observed an abrupt transition at a critical concentration around 10 nM, where condensation was observed only above this critical value (Fig. 1C). Condensation was observed for a wide range of ParB concentrations (20-200 nM) and always yielded the same degree of DNA condensation, irrespective of the ParB concentration.

Next, we optically visualized the DNA condensation in a fluorescence-based DNA-stretching assay ^35^ using a TMR-labelled *B. subtilis* ParB protein and SYTOXGreen (SxG) intercalator to label the DNA molecules. Here, the DNA*_parS_* was 42 kbp long, loosely tethered onto the surface via biotin-streptavidin interaction at both DNA ends, and it contained a single *parS* site in the middle of the DNA*_parS_* ^35^ (see Methods for details, Fig. S1D). Upon incubation with 25 nM ParB, we observed strong fluorescent DNA loci arising over time on the DNA*_parS_*, which colocalized with the fluorescent ParB signal (Fig. 1D, Fig. S1E). From the fluorescence intensity of the ParB:DNA condensate, we quantified the amount of DNA within the condensate as well as the force that the condensation exerted on the DNA fragments outside of the condensate ^36, 37^. Strikingly, we observed that DNA condensate sizes reached up to 25 kbp, i.e., it could contain over 60% of the total DNA content (see Fig. S1F). Furthermore, we observed that condensate sizes varied by a factor of >2 as the end-to-end length of the tethered DNA varied between 20% and 50% of the DNA contour length (contour length was ∼14 μm; Fig. 1E). While the condensate size thus could vary significantly, we found, by contrast, that the pulling force that it exerted on the DNA was rather constant, at a value of F = 0.08 ± 0.03 pN (mean ± SD., Fig. 1F).

Finally, we used AFM to image the ParB:DNA condensates at single-molecule resolution. The AFM images showed protein-bound DNA structures that are locally bridged and entangled, where higher local structures (>4 nm) indicated the presence of ParB molecules on the DNA (Fig. 1G-H). In contrast, we did not observe such higher-order DNA:ParB structures in the absence of either ParB, a *parS* site on the DNA, or CTP molecules (Fig. S1G-I).

### DNA condensation by ParB proteins is dynamic and reversible

Previous work on ParB proteins hinted at a dynamic exchange of the proteins with DNA*_parS_* ^24, 25^. Here, using our DNA-stretching assay, we tested the binding and unbinding of ParB within condensed ParB:DNA clusters. We co-imaged the DNA condensate and ParB proteins over time and performed fluorescence recovery after photobleaching (FRAP) experiments to assess the dynamic exchange of ParB proteins within a single cluster (Fig. 2A-B). Following bleaching, the FRAP experiments showed a quick recovery (<1 min) of the ParB fluorescence signal within the condensate (Fig. 2B), pointing to a fast exchange of ParB proteins with the soluble pool of ParB in the buffer, while maintaining a condensed ParB:DNA structure.

**Figure 2.**
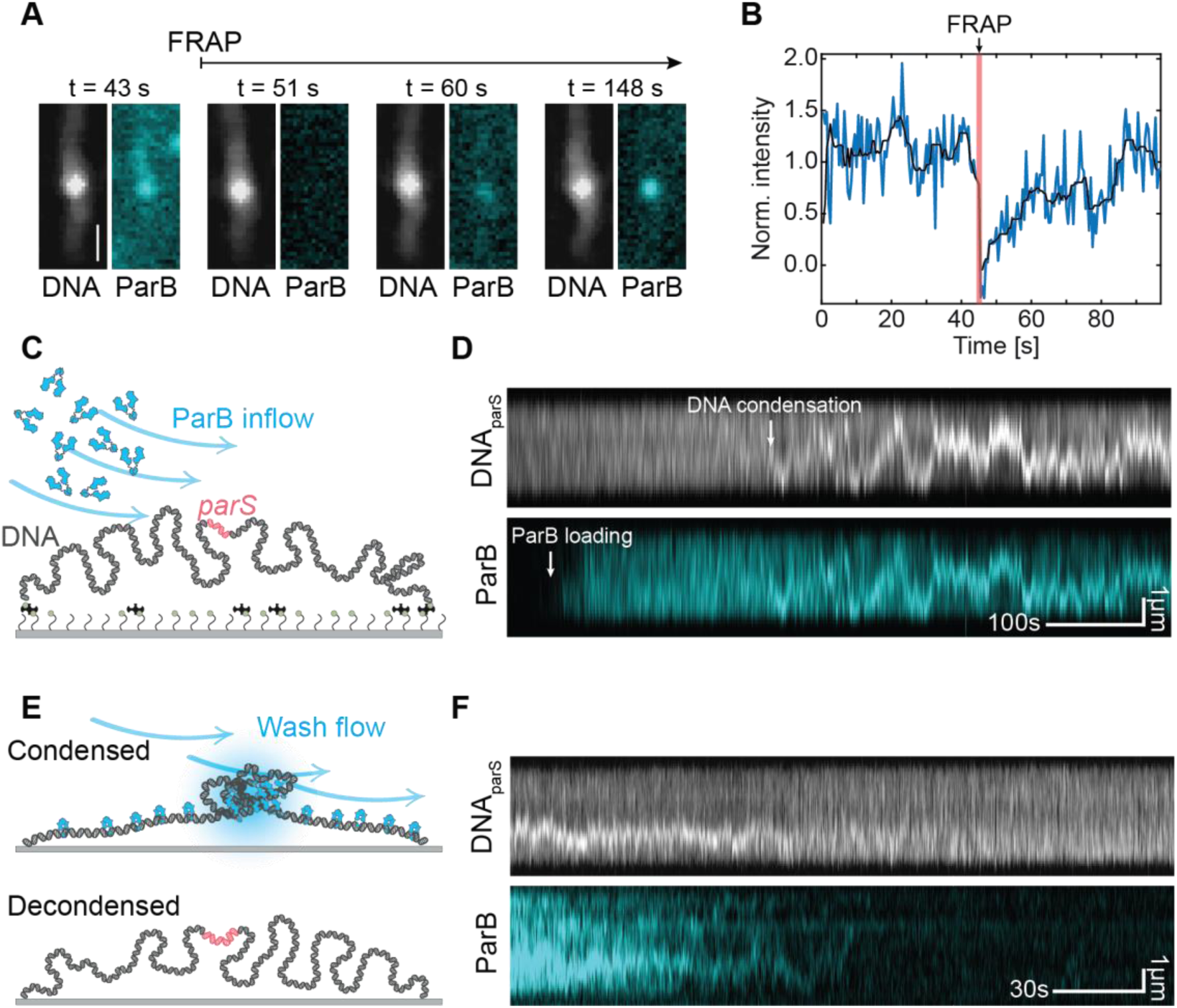
DNA condensation is dynamic. **(A)** Snapshots of DNA signal stained with SxG (greyscale) and ParB^Alexa647^ signal (cyan) shown at different time points before photobleaching and during fluorescence recovery. (**B)** Fluorescence intensity of the ParB:DNA condensate before photobleaching and during fluorescence recovery of the condensate, integrated over the full length of the DNA molecule (full trace Fig. S2A). Black line represents a median filter with a window size of 11 frames. (**C)** Schematic representation of the single-molecule fluorescence DNA-stretching assay used for probing DNA condensation. (**D)** Kymographs of DNA*_parS_*, stained with SxG (top) and ParB^TMR^ (bottom). White arrow indicates ParB loading to DNA. (**E)** Same as in C but for washing experiments. (**F)** Same as in D but for washing experiments. See also Figures S2-S3.

Interestingly, the ParB signal in the condensate did not exhibit a steady plateau but instead showed significant and sustained dynamic fluctuations of up to 40% in its intensity on the minute timescale (Fig. S2A). We quantified the total amount of DNA and ParB proteins within a single condensate over time and observed that indeed both the DNA and ParB intensities displayed substantial fluctuations, indicating that the ParB:DNA condensate varied in size over time, although the condensate was stably maintained throughout time (Fig. 2A-B and S2B-G). We recorded timelapses of the initial steps of condensation to monitor the beginning of the entire process from the ParB loading to the spreading on DNA*_parS_* and the condensation. As Fig. 2C-D shows, we observed loading of ParB, whereupon its subsequent spreading led to a covering of basically the entire DNA within minutes after flushing in the protein, after which DNA-condensation was established through the formation of a local cluster (Fig. 2D, S3A, C). The DNA*_parS_* within the resulting ParB:DNA condensates exhibited sizeable fluctuations of the order of ∼10 kbp within minutes (Fig. S3B, D). In complementary MT experiments similar high fluctuations of ParB:DNA condensates were observed (Fig. S3E, G, H). If ParB molecules would form permanent protein-protein bridges to bring distant parts of the DNA together, a continuously increasing condensation into a single static ParB:DNA condensate would occur. On the contrary, our data suggested that ParB-ParB bridges are transient, continuously reforming and breaking, thus establishing a condensate whose components are quickly turning over.

The dynamic nature of the ParB:DNA condensates was confirmed by washing experiments where we washed the condensates with a buffer that did not contain ParB proteins (Fig. 2E). Our data showed that most ParB:DNA condensates slowly washed out within a time of ∼6 minutes, wherein the DNA molecules returned to their initial non-condensed state (Fig. 2F, S4A,B). This occurred similarly with or without the presence of CTP molecules (Fig. S4A and S4C, respectively).

Altogether, FRAP experiments showed the dynamic exchange of the ParB proteins with the DNA condensate, and washing experiments pointed to the transiency of the ParB-ParB bridges that allowed for proteins to be washed away and decondense the DNA. Our data showed that the ParB:DNA condensates are dynamic entities that continuously rearrange, consistent with ParB-ParB bridges that are transient and continuously breaking and reforming while maintaining the overall condensate intact.

### DNA condensation starts with loop formation

To explore the very initial stages of the ParB-DNA condensate formation, we strongly lowered the concentration of the ParB protein to 0.3 nM and observed their behavior at the single-molecule level in real-time. Using our fluorescence DNA-stretching assay, we observed that two ParB dimers loaded on the DNA*_parS_* could spontaneously meet and connect, forming a DNA loop between them that was both dynamic in its position (as it diffused along the DNA) and transient (as it disrupted after a ∼30s) (Fig. 3A). This observed process is in line with our recently discovered ParB-ParB *in-trans* recruitment whereby a ParB dimer was shown to mediate the loading of an additional dimer at a distant genetic sequence that is proximal in the 3D space (Fig. 3B) ^35^.

**Figure 3.**
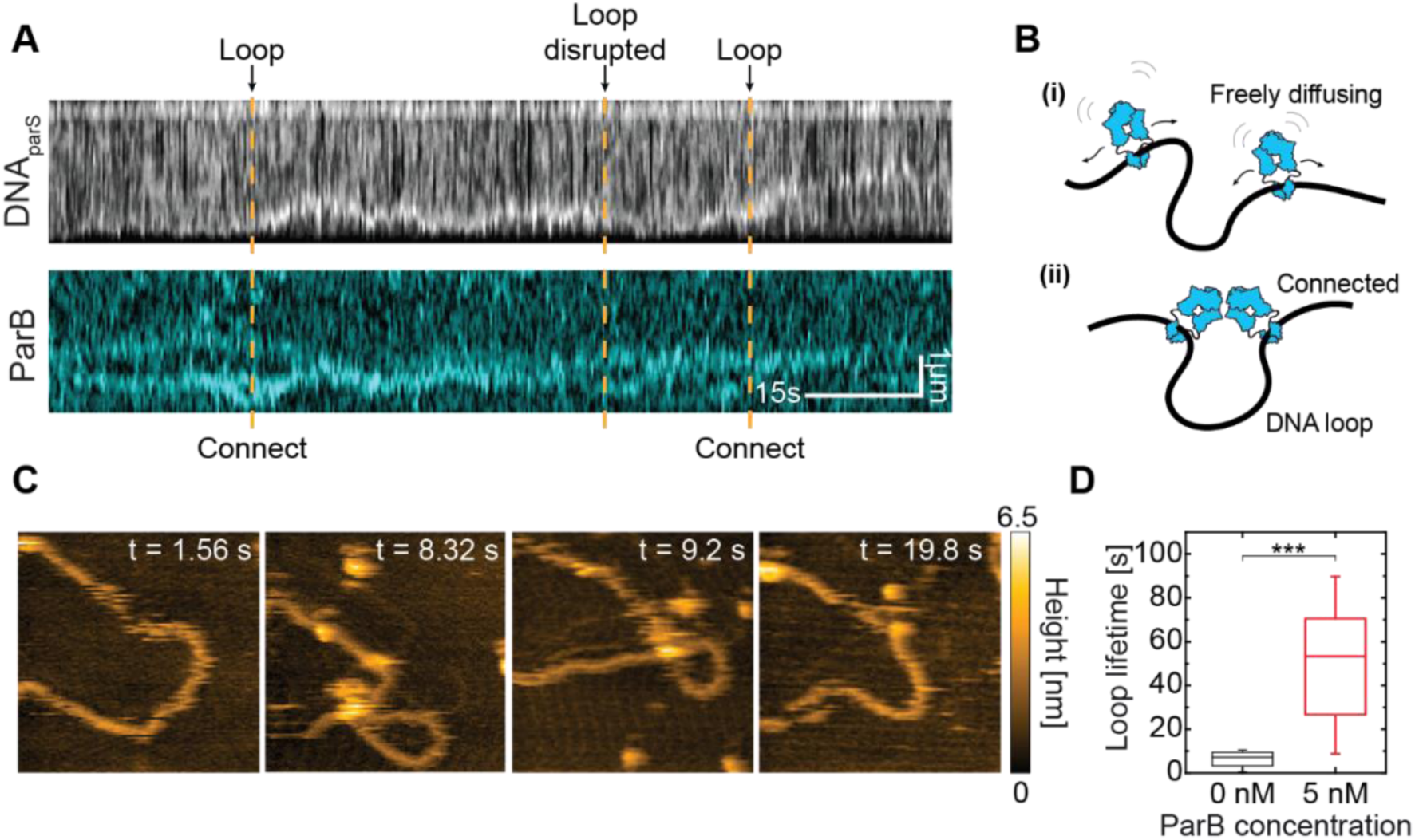
DNA condensation is initiated via the ParB-ParB bridge and a transient loop formation. **(A)** Kymographs for DNA*_parS_* stained with SxG (top) and ParB^TMR^ (bottom). Orange dashed lines show the initiation and breaking points of the DNA loop held by ParB connection points. **(B)** Schematic representation of DNA loop formation induced by ParB-ParB bridging. (**C)** High-speed AFM topography images of a transient loop formation induced by ParB-ParB bridging (see Movie S1). (**D)** Box-whisker plot of DNA loop lifetime in the absence (black; mean±SD 5.6 ± 3.3 s; N = 15) and the presence (red; mean±SD 50 ± 25 s; N = 16) of 1 nM ParB. Significance value by 2-tailed, unpaired t-test: p = 1.29 *10^-4^. See also Figure S4.

To observe the initial loop formation by ParB proteins more accurately, we turned to high-speed AFM (see Methods), where we examined a shorter circular DNA*_parS_* molecule (4.2 kbp). We observed that ParB dimers bound to different spots on the DNA could form a transient loop of DNA by bridging these distant segments together (Fig. 3C, S4D-G). We quantified the loop residence times and observed that loops persisted much longer in the presence of ParB proteins than loops observed in a control without ParB (Fig. 3D, S4H). On average, the DNA loops resided for 50 ± 25 s (mean ± SD) before disrupting. We reasoned that the observed loop formation was the precursor of the condensation of more extended DNA:ParB structures. Specifically, at larger ParB concentrations, the amount of binding ParB became larger than set by its unbinding rate, triggering a non-linear cooperative binding and spreading on the DNA substrate, leading to condensation.

### CTP hydrolysis at the ParB N-terminal is required for partition complex formation

Previous work ^6, 8, 9, 22, 23, 27, 30^ hypothesized that DNA bridging induced by ParB dimers involves the interaction between two monomers of different ParB dimers. However, current models cannot distinguish whether ParB proteins need to be opened at the N- or C-terminal domain (Fig. 4A) prior to DNA condensation^27^. Balaguer *et al*. ^27^ found that CTP binding was crucial but CTP hydrolysis was not required for DNA condensation ^27^, which suggests a third alternative where ParB dimers condense DNA without any opening after DNA loading and prior to ParB-ParB bridge formation (Fig. 4A).

**Figure 4.**
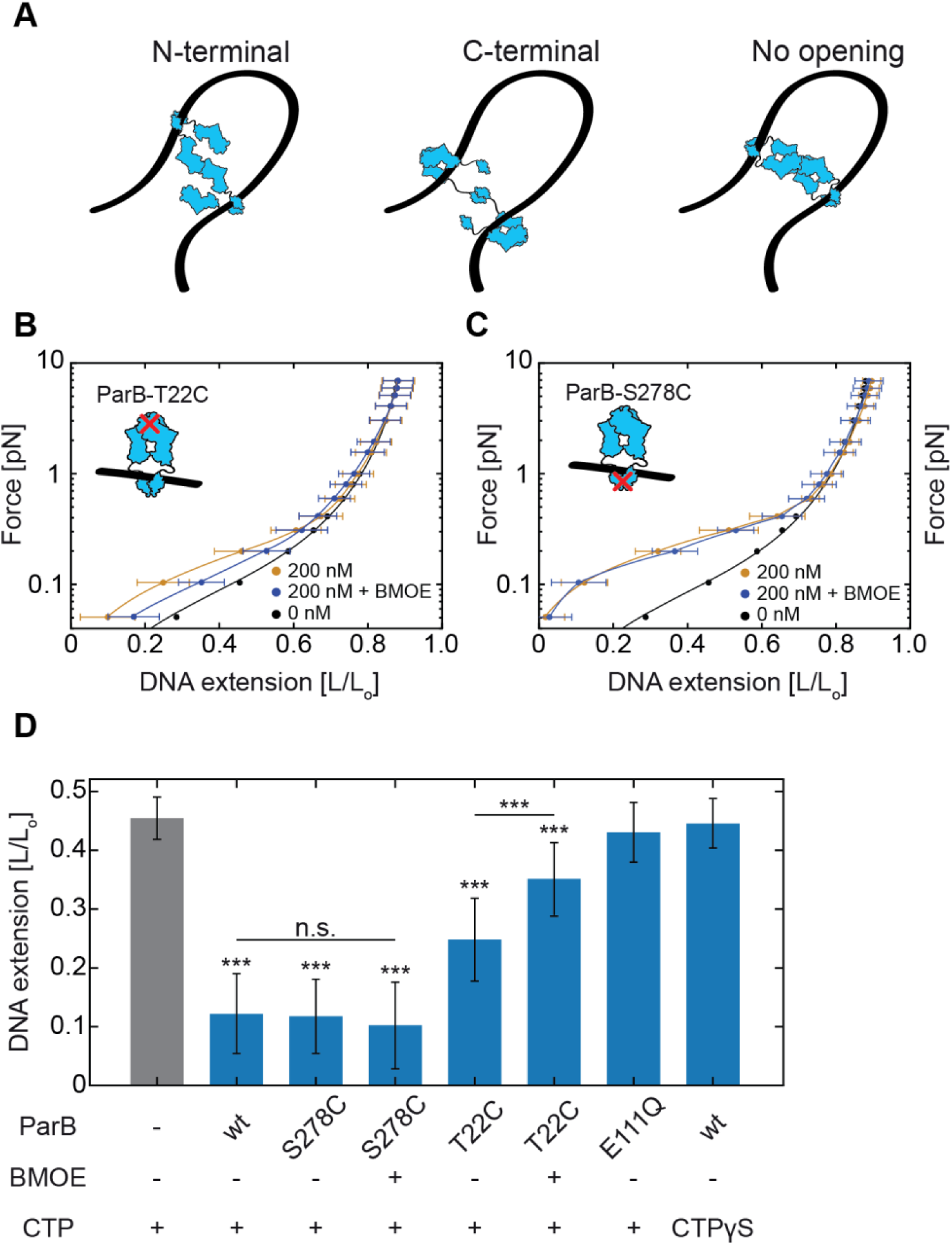
DNA*_parS_* condensation requires CTP hydrolysis at the N-terminal of ParB proteins. **(A)** Potential scenarios leading to ParB-ParB bridging *via* different protein domains. (**B)** Average DNA*_parS_* extension (mean±SD) in MT when lowering force in the presence of ParB**^T22C^**mutant with (purple; N = 36) or without (yellow; N = 43) 1 mM BMOE crosslinker added. Black line represents the WLC model fit to 0 nM ParB data (N = 32). (**C)** Same as in (B) for the C-terminal ParB**^S278C^** mutant (N; 0 nM: 32, 200 nM: 13, 200 nM +BMOE: 13). (**D)** Relative DNA extension in different conditions at an applied force of F = 0.1 pN and 200 nM concentrations of all ParB variants. Statistical analyses consisted of unpaired, two-tailed t-tests (p: *** < 0.001; n.s. = non-significant). See also Figure S5.

To distinguish between these models and resolve whether ParB needs to open after loading to DNA*_parS_* to proceed with DNA condensation, we examined a variety of ParB variants, which were forced into either N-or C-terminal domain closure. After incubating our 15 kb DNA*_parS_* with a ParB variant at a certain concentration, we collected MT force-distance curves that characterized the degree of condensation (Fig. 4B-D). Prior to spreading, ParB dimer binds as an ‘open clamp’ to the *parS*-site and sandwiches two CTP nucleotides at the N-termini interfaces that proceed to further close forming a ‘closed clamp’ ^43^. To force the protein to remain N-terminally closed after binding the DNA, we used a previously described ParB variant, namely ParB**^T22C^**, which can be efficiently crosslinked by bismaleimidoethane (BMOE) via cysteine chemistry in its closed (but not in the open) form ^28^ (Fig. S5A-B). We examined this construct in MT experiments, albeit with a different experimental procedure than previously described (Fig. 1): we first loaded the ParB**^T22C^** proteins for 5 min and then crosslinked them while bound to the DNA for 10 min, using a buffer containing BMOE and CTP but no protein (see Methods, Fig. S5A). A comparable experimental procedure increased the DNA residence time of ParB**^T22C^** in a Biolayer Interference (BLI) assay, presumably by locking ParB clamps onto DNA by cysteine cross-linking ^26, 28, 29^ (Fig. S5C-D). After loading the ParB**^T22C^** dimers on the DNA, we started the force-extension experiments (as in Fig. 1C) and observed two key differences (Fig. 4B) compared to wildtype ParB.

First, without the BMOE crosslinking, a significant condensation was observed, as can be seen from comparing the yellow curve (with ParB**^T22C^)** to the black curve (bare DNA), although the ParB**^T22C^** variant condensed the DNA to a lesser degree than wild-type ParB (Fig. 4D). Secondly, and more interestingly, the ParB**^T22C^** variant’s ability to condense the DNA was hampered in the presence of BMOE. To compare with a mutant cross-linking in the C-terminal domain^27^, we subsequently employed a ParB**^S278C^** variant that can be efficiently crosslinked by BMOE at its C-terminus while still allowing both the CTP a *parS*-site binding ^28^. The ParB**^S278C^** mutant was crosslinked prior to the experiments, ensuring that upon their contact with the DNA*_parS_*, they were already crosslinked at their C-terminus. In the MT experiments we observed no differences between the crosslinked and non-crosslinked variants (Fig. 4C)). In the presence of BMOE (and thus a covalent bond at the C-terminus), the ParB**^S278C^** proteins could efficiently load and condense the DNA*_parS_* similar to the wild-type (Fig. 4D). We confirmed the crosslinking of both the ParB**^T22C^** and ParB**^S278C^**variants by denaturing SDS-PAGE (Fig. S5A-B).

Furthermore, we examined the effects of CTP hydrolysis on ParB-mediated DNA-condensation. First, we measured the previously described ParB**^E111Q^** mutant, which is deficient in CTP hydrolysis but not in DNA*_parS_* loading ^28^. In MT force-extension experiments under the same conditions as the experiments in Fig 1C, this variant did not show any apparent DNA condensation (Fig. 4D, S5E). This is in agreement with previous studies showing that catalytically inactive EQ-mutants do not form fluorescent foci characteristic of a functional partition complex in *B. subtilis* ^28^. Finally, we performed MT experiments with replacing CTP with its non-hydrolyzable version CTPγS, which would force a clamped state of ParB after loading onto the DNA*_parS_* ^4, 29^. In line with our previous results, we observed no condensation of DNA*_parS_* molecules, even in the presence of high protein concentrations (200 nM, Fig. 4D, S5F, G).

Taken together, we can conclude that our findings demonstrate that CTP hydrolysis at the N-terminal domain is essential for DNA condensation. Most likely, this involves an N-terminal opening, while C-terminal opening is not required.

### Cooperativity of ParB multimers drives ParB:DNA condensation

We then questioned whether the interactions between ParB proteins involve merely dimer-dimer pairs or larger oligomers. In the case of “dimer of dimers” (DoD), ParB dimers behave as single bivalent bridges (Fig. 5A). By contrast, “Multimers of dimers” (MoD) can undergo oligomerization (Fig. 5A) into larger complexes. We tested these two scenarios in Molecular Dynamics (MD) simulations (Fig. 5B), where we modelled ParB dimers loading at *parS*, diffusing on a DNA substrate, and undergoing *cis/trans* recruitment (see Methods and Tišma et al. ^35^ for more details). By varying the ratio *κ* of loading versus unloading rates, we accounted for different ParB concentrations in the bulk.

**Figure 5.**
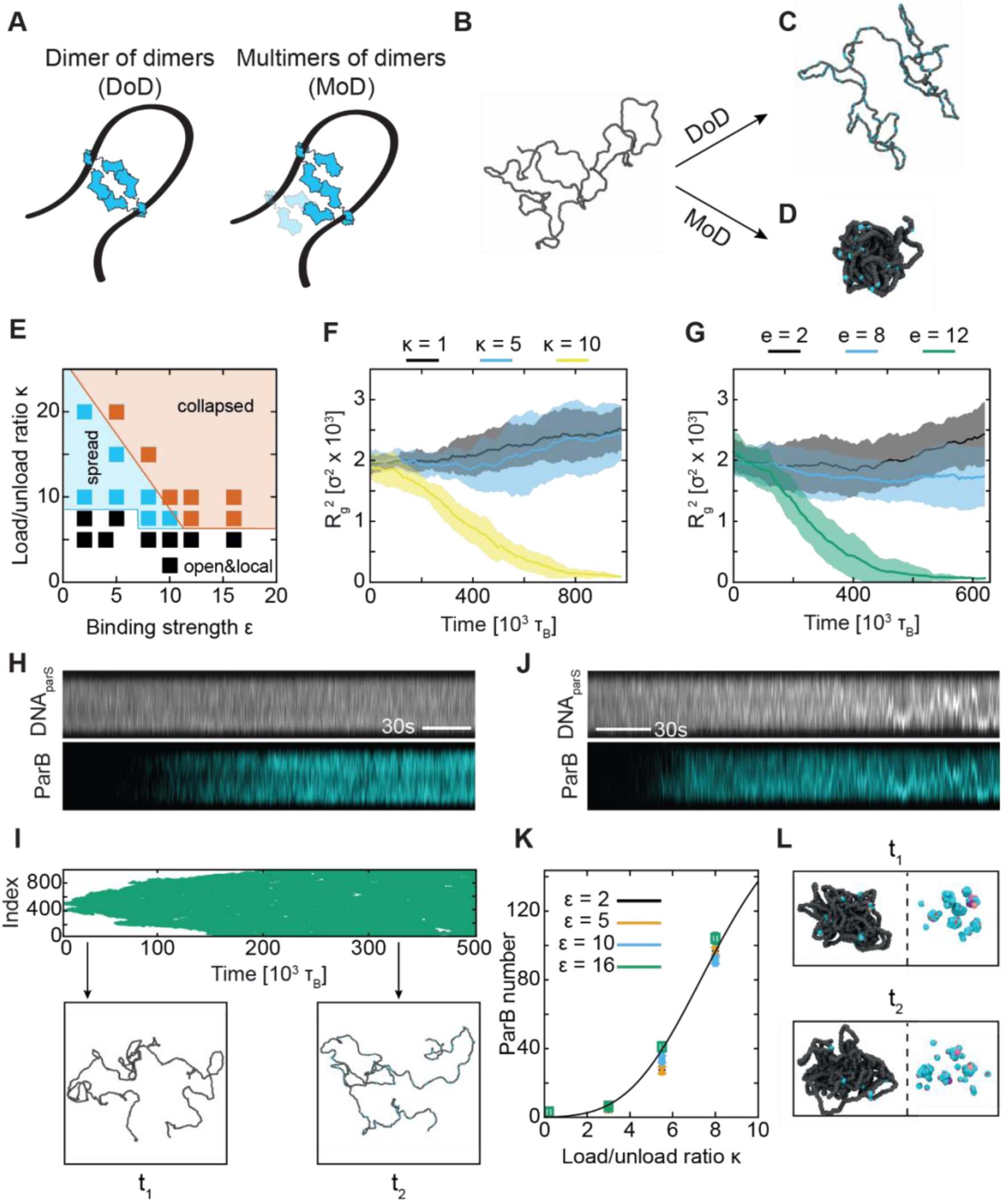
DNA cluster formation depends on bridging strength and ParB loading rate. **(A)** Two possible scenarios for ParB-ParB bridging *via* the N-termini. (**B)** MD simulations image of a DNA polymer containing 1,000 beads with non-tethered ends. (**C)** ‘Open’ DNA polymer conformation in the DoD (valency = 1) mode of ParB-ParB interaction. (**D)** Collapsed DNA polymer conformation in the MoD (valency = 2) mode of ParB-ParB interaction. (**E)** Phase diagram for the MoD model showing three distinct conformations with varying ParB-ParB interaction strength (ε) and load/unload ratios (κ). White area: local distribution of ParB and open DNA conformation; cyan area: spread distribution of ParB and open DNA conformation, and orange area: collapsed ParB:DNA conformation. (**F)** DNA collapse for varying loading/unloading ratios (with a fixed ε = 10, see Methods). R_g_^2^ represents the gyration radius of the DNA. Shaded area is SD among 10 replicas of the same system with a different initial configuration and seed. (**G)** MD simulations of DNA collapse/condensation in the presence of a fixed loading/unloading ratio (κ = 10) at varying ParB-ParB binding strengths. Shaded area: same as in F. (**H)** Kymographs for DNA*_parS_* stained with SxG (top) and ParB^TMR^ (bottom) at 5 nM. (**I)** Top: simulated kymograph of ParB distribution at ε = 5, κ = 15, where no DNA polymer collapse was observed. Bottom: snapshots of DNA polymer conformations at selected timepoints in the kymograph above. (**J)** Same as H for the ParB concentration of 25 nM. (**K)** Total number of ParB bound to the DNA as a function of κ. Black line represents fit with a Hill coefficient of H = 4.5, indicating a cooperative, switch-like behavior. (**L)** Different timepoints after condensation of ParB:DNA cluster in MD simulations (ε = 10, κ = 10). Left: shows the ParB:DNA cluster conformation. Right: shows the ParB proteins within the cluster. Three ParB proteins labelled in shades of pink to show that they dynamically rearrange while the overall DNA structure remains collapsed. See also Figure S6, S7.

For all parameters, we observed a dramatic difference in the DNA polymer structure between DoD and MoD models. When ParB dimers could only interact as DoD (valency 1), DNA molecules remained in a linearly extended, non-condensed conformation (Fig. 5C; Movie S2, S3). However, when we considered the MoD mode with valency 2, we observed the formation of a dense cluster (Fig. 5D; Movie S4) that resembled our fluorescence data (Fig. 1). Thus, for all other parameters being equal, the DoD-mode remained in fully extended conformation while MoD-mode dynamically collapsed the DNA into a condensed state.

We then extended our MD simulations to test for DNA condensation at different values of the interaction strength *ϵ* between ParB proteins in the MoD model (i.e., the depth of the Morse potential): we observed three qualitatively different outcomes (see phase diagram in Fig. 5E): (i) extended DNA with merely localized ParB dimers (Fig. 5E, white area), (ii) extended DNA where ParB proteins spread to the whole DNA substrate driving no compaction (Fig. 5E, cyan area), and (iii) condensed DNA:ParB structures (Fig. 5E, orange area). In our MD simulations, we did not observe condensation without spreading, while we did observe spreading without condensation, e.g., DNA molecules showing ParB proteins across the full length of the DNA polymer but no collapse of the polymer (Fig. S6A, B; Movie S5). This was dependent on the loading/unloading rate *κ* and interaction strength *ϵ* (Fig. S6C-E). For a sufficiently large number of ParB proteins (Fig. 5F) and ParB-ParB bridge strengths (Fig. 5G), clusters would appear (Fig. S6F, G) and the DNA molecule would collapse (Fig. 5F, G; Movie S4). This is in line with the behavior we observed in our fluorescence imaging assay: at 5 nM, we observed DNA molecules that displayed a uniform coverage of ParBs without evidence of condensation (Fig. 5H), just as in the MD simulations (Fig. 5I). On the contrary, at higher protein concentrations (25 nM), the DNA molecule collapsed shortly after ParB proteins spread over the entire DNA molecule (Fig. 5J).

This behavior can be attributed to the dynamic nature of ParB dimers on DNA. If the loading and recruitment rate (proportional to the concentration of ParB in the soluble pool) is slow compared with the unbinding time – set by the CTP hydrolysis rate^4, 28, 29^ – it is not possible to reach a critical number of ParB proteins on the DNA to drive condensation. In our MD simulations, we therefore quantified the number of ParB bound to the DNA as a function of ParB concentration (*κ*_*on*_ in our simulations, Fig. S7A-B). We obtained a steep Hill coefficient (H = 4.5, Fig. 5K) indicating high cooperativity. For instance at *κ* = 8, we observed ∼100 ParB molecules on the DNA polymer that drive the condensation (Fig. S7C). An abrupt transition was also observed in the radius of gyration of the DNA as a function of *κ*_*on*_ (Fig. 5G). This switch-like collapse behavior was also seen in both our fluorescence (Fig. 5J) and MT (Fig. 1C) experiments, where at ∼10 nM ParB the DNA molecules were either found stretched or condensed, but never in between, indicating a bi-stable nature of this cooperative, first-order-like condensation.

Finally, by visual inspection of our simulation results, we could discern that the ParB:DNA condensate was maintained by a fluctuating number of ParB proteins. Figure 5L shows example time snapshots where ParB proteins were exchanged with proteins of the pool, whilst the DNA scaffold remained condensed. This turnover that we observed in MD simulations is in line with our FRAP experiments (Fig. 2).

### Transcription by RNA Polymerase is not affected by dynamic ParB:DNA clusters

It has been reported that gene expression can be downregulated in the vicinity of *parS* sites ^38–43^. To address the long-standing question of whether ParB acts as transcriptional repressor by sterically hindering RNAP, we examined *in vitro* transcription in our single-molecule assays with DNA:ParB condensates. We included a single T7A1 promotor within the 16.5 kbp DNA_parS_ construct with a single *parS* site (Fig. 6A) together with a singly biotin-labeled bacterial RNAP from *E. coli* that efficiently binds to the T7A1 promotor and able to transcribe ∼5,000 nt long RNA constructs (Fig. 6B, D). After stalling the RNAP at position A29 after the T7A1 promoter on the DNA*_parS_* and tethering the DNA*_parS_* to the surface via DIG:anti-DIG interaction (see Methods), we incubated the flow cell with streptavidin-coated magnetic beads that bound to the stalled biotin-RNAP on the DNA*_parS_* ^44^. To ensure that a transcribing RNAP moved towards the ParB-cluster, the T7A1 promotor was located 1 kbp upstream of the *parS*. We first exposed the DNA*_parS_* to a high concentration of ParB proteins (50 nM), which ensured efficient loading and DNA condensation, as in our previous experiments (e.g., Fig. 1). Unlike the DNA section between the surface tether point and RNAP, which was held at a constant force of 5 pN, the residual 12 kbp of DNA*_parS_* experienced 0 pN force, ensuring effective DNA condensation by the high ParB concentration (cf. Fig. 1C). After the DNA*_parS_* was condensed by ParB, we initiated RNA transcription by adding 1 mM of all four NTPs (Fig. 6C).

**Figure 6.**
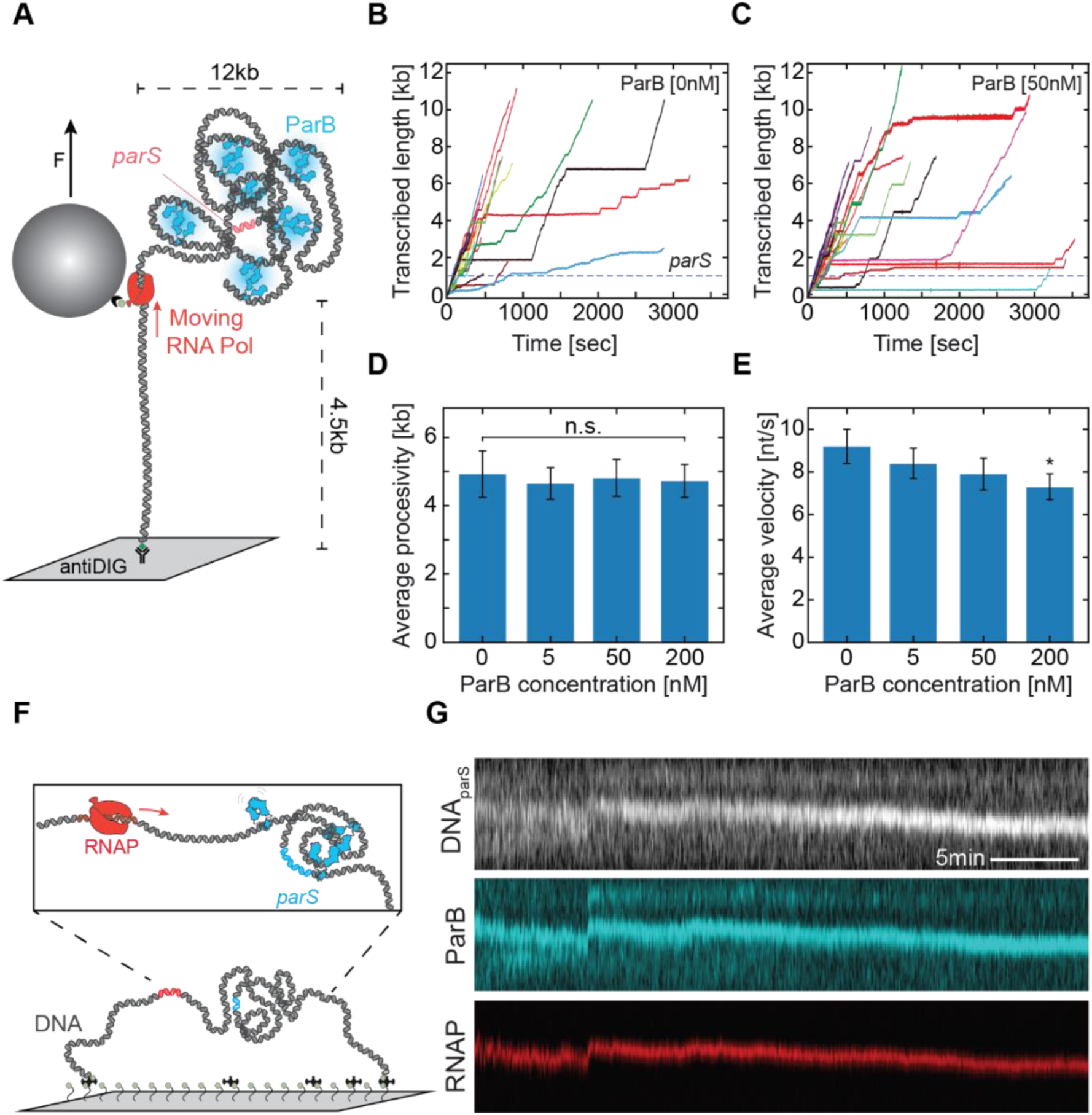
Transcription is unaffected by condensed ParB:DNA clusters. **(A)** Single-molecule MT RNAP transcription assay. (**B, C)** Individual RNAP transcription trajectories in the absence (B; N = 27) and presence (C; N = 29) of 50 nM ParB. (**D)** RNAP processivity (mean±SEM) with increasing concentrations of ParB. (**E)** Average RNAP velocity (mean±SEM) with increasing ParB concentrations (0 nM: N = 27, 5 nM: N = 46, 50 nM: N = 29, 200 nM: N = 51). Statistical analyses consisted of an unpaired, two-tailed t-test (* = p < 0.05; n.s. = non-significant). (**F)** Co-visualization single-molecule fluorescence DNA-stretching assay. **G)** Kymographs for DNA*_parS_* stained with SxG (top; grey), ParB^TMR^ (middle; cyan), and RNAP^Alexa647^ (bottom; red). See also Figure S8.

Strikingly, we observed that RNAP molecules were able to efficiently translocate along DNA despite the presence of *parS* and ParB, with virtually no effect on the average processivity or pause-free elongation rate (Fig. 6D and Fig. S8A, respectively). Even at 200 nM ParB, the processivity was not affected (Fig. 6D), indicating that a ParB:DNA cluster does not act as a transcriptional roadblock. A modest decrease ∼20% in average transcription velocity was only detectable at 200 nM ParB (from 9.2 ± 0.8 nt/s to 7.3 ± 0.6 nt/s, mean ± SEM, Fig. 6E). Notably, the decrease in average velocity resulted from the increased pausing probability while the elongation rate remained the same (Fig. S8B,A, respectively). More specifically, a probabilistic dwell time analysis of transcriptional pausing (Fig. S8D) showed that the probability of long pauses increased, whereas short elemental pausing remained constant (Fig. S8C-D)^44, 45^, and revealed that the ParB:DNA condensate induced an increase in (reversible) RNAP backtracking ^45^.

While RNA transcription was remarkably unaffected by the presence of ParB, an opposite scenario can be envisioned, i.e., that the DNA condensation breaks down during transcription as RNAP might break ParB-ParB bridges or detach ParB proteins from the DNA. To examine this, we visualized the effect of transcription on the ParB:DNA cluster in real-time using our fluorescence DNA-stretching assay (Fig. 6F). For this, we constructed a 38 kbp DNA*_parS/T7A1_* that includes a T7A1 promotor 1 kbp downstream from the *parS* site (see Methods; Fig. S8E). We used a TMR-labelled ParB (ParB^TMR^) and an Alexa647-labeled RNAP (RNAP^Alexa647^, Fig. S8F) for visualization. We observed efficient DNA condensation by 25 nM ParB in the presence of non-transcribing RNAP (Fig. S8G,H). When we subsequently initiated RNAP^Alexa647^ transcription in presence of ParB, we simultaneously observed both efficient transcription and DNA condensation (Fig. 6G), in agreement with the MT experiments. At both longer and shorter end-to-end tethering lengths, the RNAP appeared to translocate, independent of the cluster presence (Fig. 6G, S8I-L). Thus, both RNAP transcription and ParB condensation were found to be unaffected in their corresponding functions and they can simultaneously operate.

## Discussion

The ParB partition complex is a key component for the maintenance of bacterial chromosome integrity. Here, we investigated the dynamics and molecular basis of the initiation of the partition complex formation and its maintenance. Our results provide an extensive description of the dynamics of partition complex formation, the ParB domains involved in it, and the effect the partition complex on key biological functions such as transcription. Our research expands on earlier work that inadvertently often focused on nonspecific interactions in the absence of _CTP 8,9,20,21._

Using three single-molecule assays, we showed that a single *parS* site on a long DNA molecule can efficiently promote DNA condensation by ParB proteins. While previous studies using shorter DNA molecules showed a necessity for multiple *parS* sites ^27^, our MT experiments and fluorescence DNA stretching assay with long DNA molecules demonstrate that a single *parS* site is sufficient for efficient ParB loading and strong DNA condensation (Fig. 1). We hypothesize that this is due to the length of DNA molecules, where longer DNA (>10 kb) allows for spontaneous self-looping via thermal fluctuations of the polymer much more frequently than for shorter DNA molecules. Partition complex formation via a single *parS* site is in line with *in vivo* data showing that a single *parS* site continues to exhibit an unaltered phenotype in bacteria with multiple signatures of functional ParB partition complexes (i.e. fluorescence foci, SMC loading, high fidelity chromosome segregation) being preserved _18,32,46,47._

The results from our single-molecule fluorescence and MT experiments revealed that ParB and DNA form a very dynamic structure that varies in the number of ParB molecules and amount of DNA in a condensate, with sizeable fluctuations of up to 10 kbp/minute (Fig. 1, S3, S4). This dynamic rearrangement of ParB:DNA partitions is well compatible with the observed high protein turnover within the condensate and the transiency of ParB-ParB bridge formation that underlies DNA condensation (Figs. 2,3). Our FRAP recovery data also showed a high turnover of ParB proteins on the DNA molecule, while our hsAFM experiments directly observed that transient bonds can be formed between two ParB dimers to bring distal DNA segments close together (Fig. 3B-C, S6). Interestingly, the survival time of these DNA loops was roughly similar of the ParB residence time on the DNA (Fig. 3D and Ref. by Tišma et al. ^35^).

On the molecular scale, our data showed that CTP hydrolysis at the N-terminus is needed for DNA condensation. This contrasts a previous study^27^ that reported that CTP hydrolysis and clamp opening was not required for DNA condensation, which may be due to the occurrence of multiple *parS* sites in that study, where ParB proteins nucleating on the multiple loading *parS* sites may bridge and condense the DNA, or due to recently reported ParB multimers^48^ that could bridge from different *parS* sites. Our experiments using three different strategies for N-terminal closure (non-hydrolyzing protein mutant, non-hydrolyzable nucleotide, and crosslinked proteins) pointed towards a necessity for ParB opening at the N-terminus. CTP hydrolysis at the N-terminus (Fig. 4) appears to be necessary for DNA condensation *in vivo*, too ^28, 29^, as hydrolysis-deficient ParB mutants did not efficiently form a bright foci in the cell even when they were efficiently loaded onto the chromosome^28^. Our MD simulations further showed that these bridges are formed in a way that allows for local oligomerization of ParB proteins (Fig. 5A-D).

Transiency is an essential feature for the maintenance of ParB:DNA partition complexes, which was well illustrated in our experiments as well as in previous *in vivo* research on plasmid-encoded ParB proteins ^25^. Our MD simulations shed light on the cooperative and highly dynamic nature of ParB:DNA clusters. In line with computational models of generic DNA-bridging switchable proteins, we hypothesize that the high transiency yields condensates that are self-limiting in size and preferentially drive shorter-range contacts ^49, 50^. This may help the formation of the partition complex that does not include rigid structures that could hinder other cellular machineries. Previous studies showed a considerable difference in ParB residence time on the DNA after a buffer wash in presence or absence of CTP, which points to the possible CTP exchange ^28, 29^. As our previous work indicated that ParB can reside in an open state where dimer-dimer interaction can occur,^35^ the N-terminus would be freely accessible for potential CTP exchange during this time. We hypothesize that CTP molecules can be exchanged within the ParB proteins prior to bridge formation, which allows for both ParB-ParB recruitment^35^ and DNA-condensation^27^.

Notably, the stalling force that we obtained for condensate growth is very low, i.e. a force of ∼0.1 pN, which is more than two orders of magnitude lower than the stalling forces of DNA-processing motor proteins such as RNA polymerases and helicases ^51–55^ that can exert large forces up to 25 pN ^53^ and 35 pN ^51^, respectively. As the DNA machinery at such high forces will strip any ParB clamp from the DNA, a dynamic network of proteins with a high turnover ensures to maintain the ParB:DNA partition complex. Indeed, our data showed that RNAP and ParB can simultaneously perform their respective functions, transcription and DNA-condensation (Fig. 6), respectively. We speculate that the difference in dynamics between the two processes is important to enabling them to co-function. While RNAP transcription is slow (∼10 nt/s *in vitro* (this work and Janissen et al. ^44^), ∼70 nt/s *in vivo*), ParB proteins exhibit fast and dynamic loading, diffusion, and DNA condensation of up to tens of kilobases within minutes. Continuous dynamic reforming of the ParB-DNA partition complex is likely necessary for maintaining uninterrupted RNA transcription of genes in the part of the genome that is entrapped in the partition complexes. The same likely holds for the replication machinery during the replication cycle.

Previous reports on plasmid-encoded ParB proteins showed suppression of gene expression in the vicinity of the *parS* site ^38, 39, 41^, whereas more recent reports indicate uninterrupted expression of genes around *parS* sites ^30, 38, 57^. Based on our single-molecule RNAP results, we hypothesize that the reported suppression of transcription ^38–43^ could be caused by restricted excess of RNA polymerases to tightly packed ParB:DNA complexes during the initiation stage where RNA polymerase needs to access the promotor.

In light of our results, we propose a model where the DNA-ParB partition complex is initiated by DNA-ParB and ParB-ParB interactions (Fig. 7): ParB dimers load on the *parS* site by binding two CTP molecules, whereupon they diffuse away from the *parS* site as clamps (Fig. 7B) ^4, 28, 29^. During diffusion, ParB is bound to the DNA *via* its C-terminus by non-specific interactions (Fig. 7C). Upon hydrolysis of both CTP molecules, the ParB dimer opens at the N-terminus (Fig. 7D) and at this moment ^29^ it is able to connect to another ParB dimer on a distal DNA segment, consequently forming a transient DNA loop (Fig. 7E). Subsequently, ParB can form oligomers which result in the formation of larger clusters (Fig. 7F) that are stable over time even after the initial bridge disruption (Fig. 7G). Previous work on *B. subtilis* ParB diffusion suggested that upon hydrolysis there is likely a ∼15 s period of residual binding to the DNA before the dissociation occurs^35^, and thus a post-hydrolysis open-clamp ParB might still reside on the DNA for some time and connect to a second dimer (also proposed by Taylor et al., ^21^ and Fisher et al. ^20^). In the presence of a large number of ParB proteins within a partition complex (Lim et al., ^10^ and Guilhas et al. ^25^ reported ∼400-800 molecules); a continuous occurrence of ParB-ParB bridges would allow for the maintenance of the metastable partition complex. An intrinsic transiency of such bridges would allow the cellular machineries to be unaffected in their respective functions. Further *in vivo* studies of the interaction between the ParB:DNA complex and the DNA-replication/RNA-transcription machinery could further shed light on the dynamic structure of the ParB-DNA partition complex and its vital role in chromosome segregation.

**Figure 7.**
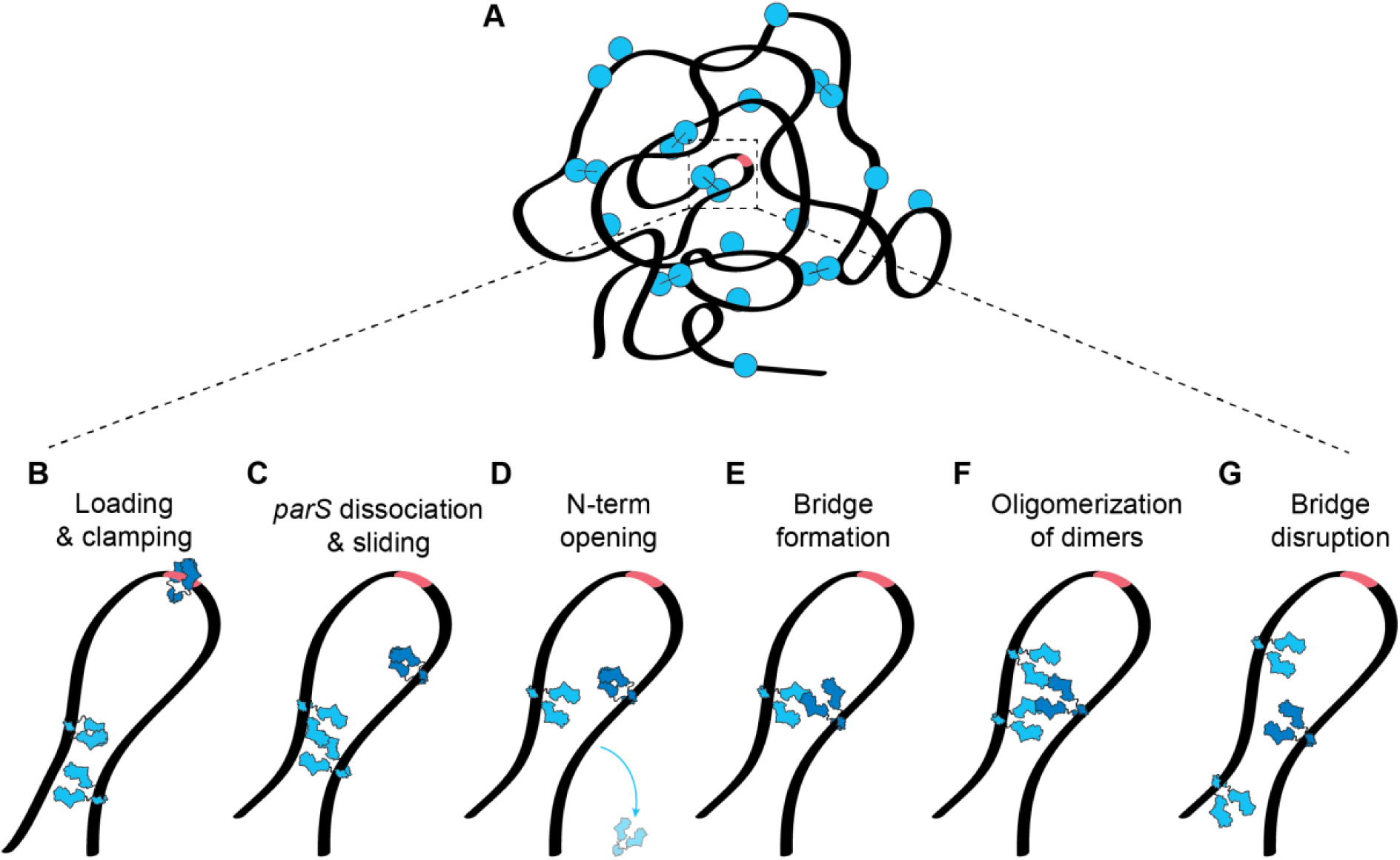
Model of dynamic ParB-ParB bridging via the N-terminal domains. **(A)** Schematic representation of a ParB:DNA cluster where few ParB-ParB bridges maintain the DNA condensate. (**B-F)** Hypothesized timeline of events experienced by ParB protein (blue). (**B)** ParB proteins load at the *parS* site and form a DNA clamp *via* dimerization at the N-terminus^4,34^. (**C)** ParB loses the affinity to *parS* site and diffuses away along the DNA ^26, 28, 29^. (**D)** After CTP hydrolysis, ParB clamp opens at the N-terminus^34^, but (**E)** remains bound to the DNA *via* its C-terminal domain^35^, to form a ParB-ParB bridge with another open clamp. This bridge is formed *via* their accessible N-termini. **(F)** Such interactions allow for oligomerization via Multimer-of-dimers mode which can stabilize the condensate over time. (**G)** After some time, a bridges can disrupt, allowing for dissociation of ParB from the DNA or connection to newly open ParB clamps, thus maintaining the fluid ParB:DNA condensate shown in panel A.

## Methods

### ParB purification and fluorescent labeling

We prepared *Bacillus subtilis* ParB expression constructs using pET-28a derived plasmids through Golden-gate cloning. We expressed untagged recombinant proteins in *E. coli* BL21-Gold (DE3) for 24 h in ZYM-5052 autoinduction medium at 24°C. Purifications of ParB and ParB^L5C^ variants, used for fluorescent labeling, were performed as described before^28^. Briefly, we pelleted the cells by centrifugation and subjected them to lysis by sonication in the Buffer A (50 mM Tris-HCl pH 7.5, 1 mM EDTA pH 8, 500 mM NaCl, 5 mM β-mercaptoethanol, 5 % (v/v) glycerol, protease inhibitor cocktail (SigmaAldrich)). We then added ammonium sulfate to the supernatant to 40% (w/v) saturation and kept stirring at 4°C for 30 min. We centrifuged the sample and collected the supernatant, and subsequently added ammonium sulfate to 50% (w/v) saturation and kept stirring at 4°C for 30 min. We collected the pellet (containing extracted ParB proteins) and dissolved it in the Buffer B (50 mM Tris-HCl pH 7.5, 1 mM EDTA pH 8 and 2 mM β-mercaptoethanol). Before loading onto a Heparin column (GE Healthcare), the sample was diluted in Buffer B to achieve the conductivity of 18 mS/cm. We used a linear gradient of Buffer B containing 1 M NaCl to elute the protein. After collecting the peak fractions, we repeated the dilution in Buffer B to 18 mS/cm conductivity and loaded it onto HiTrap SP columns (GE Healthcare). For elution, we used a linear gradient of Buffer B containing 1 M NaCl. Collected peak fractions were loaded directly onto a Superdex 200-16/600 pg column (GE Healthcare) preequilibrated in 50 mM Tris-HCl pH 7.5, 300 mM NaCl, and 1 mM TCEP. For fluorescent labeling, we incubated purified ParB^L5C^ variant with TMR-maleimide at a 1:2 molar ratio (protein:dye). We incubated the mixture for 15 min on ice, centrifuged it for 10 min, and then eluted it from a spin desalting column (Zeba) and flash frozen in liquid nitrogen. We estimated the fluorophore labelling efficiency at 70% for ParB^TMR^ monomers (90% dimers labeled) by an inbuilt function on Nanodrop using extinction coefficients of ε = 60 000 cm^-1^M^-1^.

### ParB storage and handling

We observed ParB proteins to be highly sensitive to handling and storage before the fluorescent imaging, thus handling of purified ParB protein solution is essential for high-quality experiments. Following the protein shipment between laboratories (from SG to CD lab) on dry ice, the protein solution was thawed on ice for 10 min. Concurrently, we prepared a fresh storage buffer solution 300 mM NaCl, 50 mM Tris-HCl (pH 7.5), and 1 mM TCEP and stored it on ice. We diluted the initial protein solution (∼240µM ParB) to 1.29 µM in steps of consecutive 1:1 dilutions using cold storage buffer, split the solution in 5 µl volumes and flash froze them using liquid nitrogen. On the day of the experiment, we diluted the protein once more in 1:1 ratio using a new fresh storage buffer before adding the protein at the desired concentration to either the imaging buffer in our fluorescence experiments (40 mM Tris-HCl, 65 mM KCl, 2.5 mM MgCl_2_, 1 mM CTP, 2 mM Trolox, 1 mM TCEP, 10 nM Catalase, 18.75 nM Glucose Oxidase, 30 mM Glucose, 0.25 µg/ml BSA, 25 nM SYTOX Green (SxG, ThermoFisher Scientific)) or incubation buffer in our MT experiments (40 mM Tris-HCl, 65 mM KCl, 2.5 mM MgCl_2_, 1 mM CTP, 0.25 µg/ml BSA, 1 mM TCEP). Omitting 1:1 dilutions or using buffers at different temperatures resulted in the presence of large protein aggregates in our field of view, as well as the binding of large protein aggregates to the DNA*_parS_*.

### ParBs crosslinking assays

The crosslinking experiments for ParB**^T22C^**and ParB**^S278C^** were performed under different conditions due to the following rationale: ParB**^S278C^** does not get influenced by *parS* binding because it dimerizes in solution, so the crosslinking experiment were performed and quenched before ever contacting the DNA*_parS_*, while ParB**^T22C^** needs to interact with DNA*_parS_* before closing at N-terminus binging C22 in proximity to be crosslinked, so the experiment was performed in presence of DNA*_parS_*. The crosslinking experiments were setup to mimic the MT protocol as much as possible, with obvious limitations in lack of washing steps.

We prepared the ParB**^S278C^** stock at 10 µM final concentration in the storage buffer (50 mM Tris-HCl pH 7.5, 300 mM NaCl) and kept on ice. Following this, we added BMOE to this stock to a final concentration of 1 mM, and incubated on ice for 5 min and then at room temperature for 5 min. This protein:BMOE mix was diluted 1:10 in the imaging buffer containing the quencher (50 mM Tris-HCl pH 7.5, 65 mM KCl, 2.5 mM MgCl_2_, 1 mM CTP, and 1 mM DTT) and incubated for 10 min at room temperature (equivalent to incubation/loading step in MT).

We prepared an initial loading reaction by incubating ParB**^T22C^** [2 µM] with ∼200 bp fragment containing *parS* site – DNA*_parS_* [2 µM] for 5 min at room temperature in MT imaging buffer (50 mM Tris-HCl pH 7.5, 65 mM KCl, 2.5 mM MgCl_2_, 1 mM CTP). This represents the initial loading of ParB**^T22C^** in the flow cell. Next, we added the crosslinking buffer (50 mM Tris-HCl pH 7.5, 65 mM KCl, 2.5 mM MgCl_2_, 1 mM CTP, 2 mM BMOE) in relation 1:1 and incubated for 15 min at room temperature. This represents the crosslinking via slow washing in the flow cell. We then added DTT to a final concentration of 2 mM to quench the reaction. While in our MT experiments we performed such loading and washing twice, here we could not wash away any residual components thus the final buffer contained (65 mM KCl, 50 mM Tris-HCl, 2.5 mM MgCl_2_, 1 mM CTP, 2 mM DTT, 0.5 mM BMOE leftover). We performed these experiments in multiple conditions mentioned in Fig. S5A, and such alterations to the protocol (omitting components) was performed such that the final volume remains the same, and concentration of other components remains unaltered. All conditions were ran on the 12% SDS-PAGE gel (SurePAGE™/Bis-Tris, M00668, GenScript) and the molecular weights were approximated using SeeBlue™ Plus2 Pre-stained Protein Standard (LC5925, Invitrogen).

The crosslinking efficiency was determined from a duplicate experiment (Fig. S5B) using GelAnalyzer (version 19.1).

### Biolayer interferometry assay (BLI)

We performed BLI assay as described previously by Antar et al. ^28^ (Fig. S5C-D). In brief, we used in a buffer containing 150 mM NaCl, 50 mM Tris-HCl (pH 7.5), and 2.5 mM MgCl_2_ on BLItz machine (FortéBio Sartorius). We first loaded 4 µl of 169 bp biotin-labeled DNA*_parS_* (100 nM) on the biosensor for and incubated for 5 min (Fig. S5C-D). After this loading phase, we washed the biosensor briefly once with the reaction buffer and once with the reaction buffer containing 1 mM CTP. Next, we mixed 2× ParB solution and 2× CTP solution in 1:1 ratio (final concentration of 1 μM ParB and 1 mM CTP, both in the same buffer [150 mM NaCl, 50 mM Tris-HCl (pH 7.5), and 2.5 mM MgCl_2_]), and we loaded the 4 μl of the mixture immediately on the biosensor for 2 min. In case of BMOE crosslinking test, both for ParB-wt and ParB**^T22C^**, we performed a wash using a buffer containing 1 mM BMOE [150 mM NaCl, 50 mM Tris-HCl (pH 7.5), and 2.5 mM MgCl_2_, 1 mM CTP, 1 mM BMOE] and no proteins, for 2 min. The dissociation phase was then carried for 5 min in 250 μl of protein-free reaction buffer in the absence of CTP nucleotide. We analyzed all measurements on the BLItz analysis software and replotted on GraphPad Prism for visualization.

### Construction and purification of 42 kbp DNA*_parS_* construct and 38 kbp T7A1+parS for fluorescence experiments

For the construction of a long linear DNA*_parS_*, we used a large 42 kbp cosmid-i95 reported previously ^58^ and a synthetic construct containing the *parS* site (Integrated DNA Technologies, Table. S1, underlined sequence). First, we linearized the i95 cosmid using the PsiI-v2 restriction enzyme (New England Biolabs). Next, we dephosphorylated the remaining 5’-phosphate groups using Calf Intestinal Alkaline Phosphatase for 10 min at 37°C, followed by heat inactivation for 20 min at 80°C (Quick CIP, New England Biolabs). We added the 5’-phospho group on the synthetic *parS* fragment by adding a T4 kinase for 30 min at 37°C and heat-inactivated 20 min at 65°C in 1x PNK buffer supplemented with 1 mM ATP (T4 PNK, New England Biolabs). Next, we ligated the two fragments together using a T4 DNA ligase in T4 ligase buffer (New England Biolabs), containing 1 mM ATP overnight at 16°C. The final cosmid construct was transformed into *E. coli* NEB10beta cells (New England Biolabs), and we verified the presence of insert by sequencing using JT138 and JT139 (Table S1).

The final 38kb T7A1+*parS* construct was made from plasmid 189-pBS-T7A1-parS (Table S1). This plasmid was generated from plasmids 179, 66, 67, 69, 163 and 71 using a BsaI-HFv2 golden gate reaction mix (NEB E1601). The intermediate vectors were made using traditional cloning techniques (in details described in Table S1).

To prepare a linear fragment adapted for flow cell experiments, we isolated cosmid-i95 or pBS-T7A1-parS *via* a midiprep using a Qiafilater plasmid midi kit (Qiagen). We digested the cosmid-i95 for 2 h at 37°C using SpeI-HF restriction enzyme (New England Biolabs) and heat-inactivated for 20 min at 80°C. pBS-T7A1-parS was digested with NotI-HF and XhoI under the same conditions and heat-inactivated using the same procedure. Next, we constructed the 5’-biotin handles by a PCR from a pBluescript SK+ (Stratagene) using 5’-biotin primers JT337 and JT338 (Table S1), to get a final 1,246 bp fragment. We digested the PCR fragment using the same procedure described for cosmid-i95, resulting in ∼600 bp 5’-biotin-handles. The handles for the 38 kbp T7A1-parS construct were made with the same primers, but on template plasmid 186 (Table S1, S2) resulting in a final fragment of 514 bp. We further digested these PCR fragments were digested using NotI-HF or XhoI for 2 h at 37°C. This resulted in the handles of ∼250 bp in length.

Finally, we mixed the digested constructs and handles in a 1:10 molar ratio and ligated them together using T4 DNA ligase in T4 ligase buffer (New England Biolabs) at 16°C overnight, which was subsequently heat-inactivated for 10 min at 65°C. The 38 kbp T7A1-parS + handle construct were subsequently digested for 1 h at 37°C and heat-inactivated for 20 min at 65°C using SrfI restriction enzyme (New England Biolabs).

We cleaned up the resulting linear (42 +1.2) or (38 + 0.5) kbp DNA*_parS_* constructs from the access handles using an ÄKTA Start (Cytiva), with a homemade gel filtration column containing 46 ml of Sephacryl S-1000 SF gel filtration media, run with TE + 150 mM NaCl buffer at 0.5 ml/min. We stored the collected fractions as aliquots after snap-freezing them by submerging them in liquid nitrogen.

### Single-molecule visualization assay

The surface of imaging coverslips was prepared as previously described ^59^, with the addition of surfaces being pegylated 5x 24 h. For immobilization of 42 kbp DNA*_parS_*, we introduced 50 μl of ∼1 pM of 5’-biotinylated-DNA*_parS_* molecules at a flow rate of 1 – 4 μl/min, depending on the desired end-to-end length in the experiment, in T20 buffer (40 mM Tris-HCl (pH 7.5), 20 mM NaCl, 25 nM SxG). Immediately after the flow, we further flowed 100 μl of the wash buffer (40 mM Tris–HCl, pH 7.5, 20 mM NaCl, 65 mM KCl, 25 nM SxG) at the same flow rate to ensure stretching and tethering of the other end of the DNA to the surface. By adjusting the flow, we obtained a stretch of around 15−40% of the contour length of DNA. Next, we flowed in the imaging buffer (40 mM Tris-HCl, 2 mM Trolox, 1 mM TCEP, 10 nM Catalase, 18.75 nM Glucose Oxidase, 30 mM Glucose, 2.5 mM MgCl_2_, 65 mM KCl, 0.25 µg/ml BSA, 1 mM CTP, 25 nM SxG) without ParB protein at the low flow rate (0.2 µl/min) to enable minimal disturbances to the DNA molecules before and after protein addition. Experiments were performed in the same conditions with the exception of replacing 1 mM CTP with 1 mM CTPγS where mentioned. Real-time observation of ParB diffusion was carried out by introducing ParB (5-10 nM) in the imaging buffer.

For the RNAP transcription experiments, we prepared the RNAP ternary complex as described previously by Janissen et al. ^44, 45^. RNAP holoenzyme was stalled on the DNA*_parS/T7A1_* constructs at position A29 after the T7A1 promoter sequence. To do so, we added 3 nM of RNAP holoenzyme to 3 nM linear DNA*_parS/T7A1_* template in 20 mM Tris, 100 mM KCl, 10 mM MgCl_2_, 0.05% (v/v) Tween 20 (SigmaAldrich) and 40 mg/mL BSA (New England Biolabs), pH 7.9, and incubated 10 min at 37°C. Afterwards, we added 50 μM ATP, CTP, GTP (GE Healthcare Europe), and 100 μM ApU (IBA Lifesciences GmbH) to the solution and incubated for 10 min at 30°C. To ensure that we measured the transcription dynamics of single RNAp ternary complexes, we sequestered free RNAP and RNAP that were weakly associated with the DNA by adding 100 µg/ml heparin and incubating for 10 min at 30°C. The ternary complex solution was then diluted to a final concentration of 100 pM of the RNAP:DNA*_parS/T7A1_* complex. The complex was flushed into the flow cell at the speed of 4 µl/min and subsequently washed for all unbound molecules in the buffer containing 40 mM Tris-HCl, 2 mM Trolox, 30 mM Glucose, 4 mM MgCl_2_, 70 mM KCl, 0.25 µg/ml BSA, 1 mM CTP, 25 nM SxG. After the washing of unbound RNAP:DNA*_parS/T7A1_* complexes, we first subjected the buffer to the imaging buffer without ParB proteins and lacking all NTPs (40 mM Tris-HCl, 2 mM Trolox, 1 mM TCEP, 10 nM Catalase, 18.75 nM Glucose Oxidase, 30 mM Glucose, 4 mM MgCl_2_, 70 mM KCl, 0.25 µg/ml BSA, 1 mM CTP, 25 nM SxG), for 3 min. This was followed by the same buffer with the addition of 25 nM ParB proteins (10 nM ParB^TMR^ and 15 nM WT ParB), which we incubated for 5 min to allow for DNA condensation to start prior to RNAP transcription. Following this incubation we added the final imaging buffer which allows DNA condensation and RNAP transcription (40 mM Tris-HCl, 2 mM Trolox, 1 mM TCEP, 10 nM Catalase, 18.75 nM Glucose Oxidase, 30 mM Glucose, 4 mM MgCl_2_, 70 mM KCl, 0.25 µg/ml BSA, 25 nM SxG, 1 mM NTP and 2 µM GreB) and started the imaging. In these experiments the signals were obtained by alternate excitation with 100 ms exposure times for DNA-SxG (488 nm laser), ParB^TMR^ (561 nm laser), and RNAP^Alexa647^ (647 nm laser) followed by a 2700 ms pause before the next frame. We observed 40% of the RNAP^Alexa647^ molecules undergo transcription in the absence of ParB molecules (N = 18/44, Fig. S8F shows an example trace).

We used a home-built objective-TIRF microscope to achieve fluorescence imaging. We used alternating excitation of 488 nm (0.1 mW), 561 nm (12 mW) and 647 nm (12 mW) lasers in Highly Inclined and Laminated Optical sheet (HiLo) microscopy mode, to image SxG-stained DNA and TMR-labelled ParB and Alexa647-RNAP respectively. All images were acquired with an PrimeBSI sCMOS camera at an exposure time of 100 ms, with a 60x oil immersion, 1.49 NA objective (CFI APO TIRF, Nikon).

### Fluorescence recovery after photobleaching experiments

Our FRAP experiments were performed using the same 42 kbp DNA*_parS_* mentioned above. For these experiments, we used Alexa647-labelled ParB proteins and SxG-labelled DNA, using the same experimental procedure and buffers mentioned above. The final ParB^Alexa647^ concentration was 25 nM to ensure strong DNA condensation in out fluorescence assay.

In these experiments the signals were obtained by alternate excitation with 200 ms exposure times for DNA-SxG (488 nm laser), ParB^Alexa647^ (647 nm laser). The photobleaching was performed using 100 mW laser power (647 nm) for 100 bursts. We used a home-built objective-TIRF microscope to achieve fluorescence imaging. We used alternating excitation of 488 nm (0.5 mW), and 647 nm (15 mW) lasers in HiLo microscopy mode, to image SxG-stained DNA and TMR-labeled ParB. All images were acquired with iXon Ultra 897 EMCCD (X-7105) camera 100x oil immersion objective (X-Apo, 1.45NA, Olympus™).

### Image processing and analysis

The areas with single DNA molecules were cropped from the raw image sequences and analyzed separately with custom-written, interactive python software ^60^. The images were smoothened using a median filter with a window size of 10 pixels, and the background was subtracted with the “white_tophat” operation provided in the *scipy* python module. The contrast of the images was further adjusted manually for visualization only (i.e. Fig. 2). The ends of a DNA were manually marked. The distance between DNA ends is the end-to-end length *R* of the DNA molecule. Total fluorescence intensity of 11 pixels (∼1.3 µm) across the axis of the DNA was obtained for each time point and was stacked to obtain a kymograph (i.e. Fig. 2). The same image area was selected to obtain kymograph of the ParB fluorescence channel.

DNA condensates appear as spots of the increased fluorescence signal (i.e., Fig. 1D, 2A) compared to the surrounding fluorescence signal along the DNA. The position of the spots was identified and tracked using the *trackpy* python module (v0.4.2; 10.5281/zenodo.7670439). The amount of DNA contained within the condensate is computed as extensively described previously by Davidson et al. ^61^. In brief, the condensate size is expressed as the fraction of intensity at the condensate position, *I*_*cond*_, in relation to the total intensity along the entire DNA molecule, *I*, and the DNA length *L* = 42.5 *kbp*

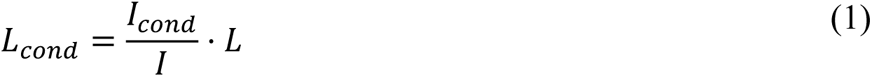

The tension acting on the DNA is expressed in terms of the contour length of the DNA outside the condensate (in bp), *L*_*out*_ = *L* − *L*_*cond*_, which is 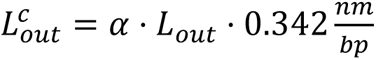, accounting for the distance between base pairs of 0.324 nm and *α* is a correction factor accounting for the lengthening of the dsDNA contour length in the presence of intercalating dyes (for 25 nM SxG, *α* = 1.0357 was used ^61^). The relative extension of the DNA outside the condensate, 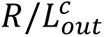 is used to compute the momentary tension acting on the DNA via the empirically determined and well-established force-extension relationship ^62^:

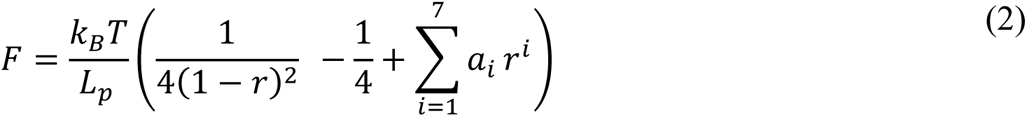

where *k*_*B*_*T* = 1.3806503 ⋅ 10^−23^ ⋅ 293 *K* ⋅ 10^−18^ *pN* ⋅ *μm*, *L*_*p*_ = 39.7 *nm* is the persistence length of DNA at 25 nM SxG, *a*_1_ = 1, *a*_2_ = −0.5164228, *a*_3_ = −2.737418, *a*_4_ = 16.07497, *a*_5_ = −38.87607, *a*_6_ = 39.49944, *a*_7_ = −14.17718.

### Construction of DNA*_parS_* construct for atomic force microscopy experiments

To construct the circular DNA for AFM experiments we used a commercially available pGGA plasmid backbone (New England Biolabs). We linearized the plasmid using MT032 and MT033 primers (Table S2). In parallel to this we extracted a region containing *parS* site downstream of *metS* gene in *B. subtilis* genome by a colony PCR using primers MT039 and MT040 (Table S2). We combined the plasmid backbone with the colony PCR insert by mixing them in molar ration 1:3 in the 2xHiFi mix (New England Biolabs). We incubated the reaction at 50°C for 60 min, and cooled down to 12°C for 30 min. We then transformed 2 µl of this reaction into the *E. coli* NEB5alpha cells (New England Biolabs), and we verified the presence of insert in grown colonies the following day by sequencing using MT030 and MT031 (Table S2). We grew sequence-positive clones for the plasmid extraction at 37°C overnight in presence of a selective antibiotic Chloramphenicol (30 µg/ml) and isolated the final plasmid using a QIAprep Spin Miniprep kit (Qiagen) the following day. The plasmids were nicked using a standard protocol for Nb. BbvCI nicking enzyme (New England Biolabs) capitalizing on a pre-existing recognition site in *metS* gene. This was done in order to remove any supercoiling from the isolated plasmids, which would alter the experimental conditions and conclusions via promoting in-trans interactions between different plasmid domains. We ligated the nicked plasmids using a T4 DNA ligase in T4 ligase buffer (New England Biolabs), containing 1 mM ATP overnight at 16°C. The following day we performed three more rounds of plasmid clean-up using the same QIAprep Spin Miniprep kit (Qiagen) to remove all the residual enzymes which could corrupt the AFM images.

### Atomic force microscopy experiments and imaging

We obtained images in dry conditions using an AFM from Bruker (Massachusetts, USA) and Scanassyst-Air-HR tips from Bruker. AFM was operated using peakforce-tapping mode. We used WSxM ^63^ software for all image processing and data extraction from our raw data in AFM experiments.

We incubated samples with different molarity ratios of DNA, CTP and ParB in Eppendorf tubes for 2 to 5 min in a buffered solution (40 mM Tris, pH 7.5, 65 mM KCl, 7.5 mM MgCl_2_). Then, we deposited the solution onto a freshly cleaved mica for 30 s. Afterwards, we thoroughly washed the surface with 3 ml of Milli-Q water and dried it under a flow of Nitrogen until visibly dry.

We took images in aqueous conditions using a high-speed AFM from RIBM (Tsukuba, Japan) and PRE-USC-F1.2-k0.15 tips from Nanotools. Here, we operated the AFM using tapping mode. The samples with DNA, CTP, and ParB were prepared in a buffered solution (40 mM Tris, pH 7.5, 65 mM KCl, 10 mM NiCl_2_) before depositing them onto a freshly cleaved and pre-incubated with Poly-L-lysine mica. Then, we thoroughly washed the surface 3 times with 8 µl of the same buffer.

### Construction of DNA*_parS_* construct for Magnetic Tweezers experiments

For the construction of a 14 kbp long linear DNA*_parS_* for MT experiments, we used a #126-pSC-T7A1reverse-parS plasmid (Table S1). Due to the plasmid size (∼20 kbp) we linearized it using 3xPCR reactions using primers MT234/MT237, MT235/MT238 and MT236/MT239 (Table S2). Concurrent with being used in linearization, MT237 primer contains an internal *parS* site - TGTTCCACGTGTAACA (Table S2). We visualized the three obtained fragments of sizes 7,783 bp, 7,638 bp and 5,594 bp for the corrected size using EtBr-agarose gel (1xTAE buffer (ThermoFisher Scientific, J63931.K2), 0.8% agarose (Promega, V3125), 0.5 µg/ml etidium-bromide (Sigma Aldrich, 1239-45-8), and extracted each from the gel using a standard protocol from PCR purification kit (Promega, A9282). We then mixed the three fragments together in 10 µl in molar ratio 1:2:1, respectively, and added to 10 µl of 2xHiFi mix (New England Biolabs). We incubated the reaction at 50°C for 60 min, and cooled down to 12°C for 30 min. We transformed 2 µl of this reaction into the *E. coli* NEB5alpha cells (New England Biolabs), and we verified the presence of desired insert (*parS* site) by sequencing using MT240 and MT241 (Table S2). We grew sequence-positive clones for the plasmid extraction at 30°C overnight in presence of a selective antibiotic Ampicillin (100 µg/ml) and isolated the final plasmid *via* a Midiprep using a Qiafilater plasmid midi kit (Qiagen) the following day.

To prepare linear fragment adapted for Magnetic Tweezer experiments, we digested #126-pSC-T7A1reverse-parS with SpeI-HF and BamHI-HF 1 h at 37°C and heat-inactivated for 20 min at 80°C (New England Biolabs). Subsequently we ran the digested fragment on a 1% TAE agarose gel and the desired ∼14 kbp DNA fragment was isolated from an agarose gel using a gel purification kit (Promega, A9282).

We made two different construct and therefore we made three different handles. For the DNA condensation experiments (e.g., Fig. 1A-C, Fig. 4), we used DNA molecules with two attachment points; Digoxigenin handle at the surface side and Biotin handle towards the magnetic bead. We made both handles using primers CD21 and CD22 (Table S2) in a PCR on pBlueScriptSKII SK+ (Stratagene). This was done in the presence of 1/5 biotin-16-dUTP/dTTP or digoxigenin-11-dUTP/dTTP (two separate reactions) resulting in a 1246 bp fragment with Biotin or Digoxigenin at different ends (JenaBioscience, NU-803-DIGXS, NU-803-BIO16-L). We digested the Biotin PCR-fragment with BamHI-HF, and the Digoxigenin-PCR-fragment with SpeI-HF, both for 2 h at 37°C. This resulted in handles of ∼600 bp in length with multi-biotin or multi-digoxigenin nucleotides present. Subsequently, for the condensation MT experiments a torsionally constrained construct was made by ligating, using T4 DNA ligase (NEB), the 14 kbp DNA fragment to multi-Biotin and multi-Digoxigenin handle in a 1:10 ratio.

For the experiments using RNAP we used a singly tethered DNA construct that only contained Digoxigenin handle at one end (attaching to the surface) while the second attachment point was on the RNAP (Fig. 6A). Thus, the second handle was a blunt end handle that did not attach to anything (Fig. 6A). Here, we made handles using the same primers CD21 and CD22 in a PCR on pBlueScriptSKII SK+ (Stratagene) either in the presence of digoxigenin-11-dUTP/dTTP, as previously, or no modified nucleotides (for blunt end side). This resulted in the same 1,246 bp fragment with Digoxigenin at one end and a second blunt end. We digested the no modification PCR-fragment with BamHI-HF, and the Digoxigenin-PCR-fragment with SpeI-HF, both for 2 h at 37°C. This resulted in the same length handles as previously (∼600 bp) with blunt end side and a multi-digoxigenin nucleotides present at the other side. For the RNAP tweezer experiments a construct with on one side multi-digoxigenin was made by ligating the 14 kbp DNA fragment to multi-Digoxigenin and unmodified handle in a 1:10 ratio. For both constructs, we cleaned up the resulting tweezer construct (14+1.2 kbp) from the access handles by running this on a 1% agarose gel and gel purify the right DNA fragment with a gel purification kit (Promega, A9282).

### *E. coli* RNAP-biotin, GreB, and σ^70^ purification

Wild-type *E. coli* RNA polymerase (RNAP) holoenzyme (α2ββ′ω) with the transcription initiation factor σ^70^ was expressed and purified as described previously ^64^. The RNAP contained a biotin-modification at the ß’-subunit ^65^ that served as an anchor to attach streptavidin-coated magnetic beads. GreB factor was obtained and purified following a previously established protocol ^66^. The activity of all purified proteins was confirmed using standard bulk transcription assays.

### Magnetic tweezers instrument and experiments

The MT implementation used in this study has been described previously^44, 45^. In short, light transmitted through the sample was collected by a 50x oil-immersion objective (CFI Plan 50XH, Achromat; 50x; NA = 0.9, Nikon) and projected onto a 4-megapixel CMOS camera (#4M60, Falcon2; Teledyne Dalsa) with a sampling frequency of 50 Hz. The applied magnetic field was generated by a pair of vertically aligned permanent neodymium-iron-boron magnets (Supermagnete GmbH, Germany) separated by a distance of 1 mm, and suspended on a motorized stage (#M-126.PD2, Physik Instrumente) above the flow cell. Additionally, the magnet pair can be rotated around the illumination axis by an applied DC servo step motor (C-150.PD; Physik Instrumente). Image processing of the collected light allows to track the real-time position of both surface attached reference beads and superparamagnetic beads coupled to DNA*_parS_* or RNAP in three dimensions over time. The bead x, y, z position tracking was achieved using a cross-correlation algorithm realized with custom-written software in LabView (2011, National Instruments Corporation)^67^. The software determined the bead positions with spectral corrections to correct for camera blur and aliasing.

The flow cell preparation for the MT experiments used in this study has been described in detail by Janissen et al. ^44, 45^. Briefly, polystyrene reference beads (Polysciences Europe) of 1.5 µm in diameter were diluted 1:1500 in PBS buffer (pH 7.4) and adhered to the KOH treated surface of the flow cell channel. Next, 0.5 mg/mL digoxigenin antibody Fab fragments (Roche Diagnostics) were incubated for 1 h within the flow cell channel, following incubation for 2 h of 10 mg/mL BSA (New England Biolabs) diluted in Buffer A containing 20 mM Tris-HCl pH 7.9, 100 mM KCl, 10 mM MgCl_2_, 0.05% (v/v) Tween 20 (SigmaAldrich) and 40 mg/mL BSA (New England Biolabs).

For the force-extension experiments, 1 pM of the 15 kbp linear DNA_parS_ was incubated in PBS buffer for 20 min in the flow cell channel. After washing with 500 ml PBS buffer, the addition of 100 µl streptavidin-coated superparamagnetic beads (diluted 1:400 in PBS buffer; MyOne #65601 Dynabeads, Invitrogen/Life Technologies) with a diameter of 1 µm for 5 min resulted in the attachment of the beads to biotinylated DNA*_parS_*; non-attached beads were then washed out with PBS buffer. Prior to conducting the force-extension experiments, the DNA*_parS_* tethers were assessed by applying a high force (7 pN) and 30 rotations to each direction. Only DNA tethers with singly bound DNA*_parS_* and correct end-to-end lengths were used for the single-molecule experiments. In the force-extension experiments, the ParB proteins (or ParB variants; see main text) were induced to the flow cell channel while applying 7 pN to the DNA*_parS_* tethers. For wild type ParB, ParB**^E111Q^**and ParB**^S278C^** we first incubated the proteins for 15 min in the flow cell at such high force and directly after, the force was gradually reduced in steps (see Fig. 1B) from 7 pN to 0.05 pN within the 15 minutes the bead z-positions were recorded. For the ParB**^T22C^** variant, we first incubated the proteins for 5 min without the presence of BMOE (65mM KCl, 50mM Tris-HCl, 2.5mM MgCl_2_, 1mM CTP) and then crosslinked them using a buffer containing 1 mM BMOE, but not ParB**^T22C^** for 10 min slow wash (65mM KCl, 50mM Tris-HCl, 2.5mM MgCl_2_, 1mM CTP, 1 mM BMOE). We repeated this step twice to ensure high presence of ParB**^T22C^** on the DNA*_parS_*. Following these two rounds of loading and crosslinking ParB**^T22C^** we performed a washing step in the buffer containing no ParB**^T22C^** proteins and a quencher (65mM KCl, 50mM Tris-HCl, 2.5mM MgCl_2_, 1mM CTP, 2 mM DTT), to ensure that any residual BMOE is quenched, while any non-crosslinked proteins are washed away before the force-extension experiments began. After this the force was gradually reduced and the z-positions of the beads were recorded, as mentioned above.

For the RNAP transcription experiments, the preparation of the RNAP ternary complex was performed as described previously^44, 45^. Briefly, RNAP holoenzyme was stalled on the DNA*_parS/T7A1_* constructs at position A29 after the T7A1 promoter sequence. To do so, 3 nM of RNAP holoenzyme was added to 3 nM linear DNA*_parS/T7A1_* template in Buffer A and incubated 10 min at 37°C. Afterwards, 50 μM ATP, CTP, GTP (GE Healthcare Europe), and 100 μM ApU (IBA Lifesciences GmbH) were added to the solution and incubated for 10 min at 30°C. To ensure that we measured the transcription dynamics of single RNAP ternary complexes, we sequestered free RNAP and RNAP that were weakly associated with the DNA by adding 100 µg/ml heparin and incubating for 10 min at 30°C. The ternary complex solution was then diluted to a final concentration of 250 pM of the RNAP:DNA*_parS/T7A1_* complex. The complex was flushed into the flow cell and incubated for 30 min at room temperature. The subsequent addition of 100 μl streptavidin-coated superparamagnetic beads (diluted 1:400 in buffer A; MyOne #65601 Dynabeads, Invitrogen/Life Technologies) with a diameter of 1 μm resulted in the attachment of the beads to the stalled biotinylated RNAP. With the introduction of ParB proteins of different concentrations (see concentrations used main text), transcription was re-initiated by added ATP, CTP, GTP, and UTP (GE Healthcare Europe) at equimolar concentration of 1 mM and immediately starting the single-molecule measurements. The experiments were conducted for 2 h at constant force of 5 pN.

All MT data sets were processed and analyzed using custom-written Igor v6.37-based scripts ^45^. From our raw data, we removed traces showing surface-adhered magnetic beads as well as tethers in our force-extension experiments where the DNA-bead attachment points were far from the magnetic equator of the beads using a previously described method ^68^. Tethers that detached from the surface during the measurement were also rejected from further analysis. All traces resulting from experiments conducted at identical conditions were pooled. The absolute z-position of the RNAP during the transcription process was converted to transcribed RNA product as a function of time, using the end-to-end length determined by the extensible worm-like chain model ^69^. The extensible worm-like chain model was also used for the fitting to the force-extension DNA_parS_ end-to-end length data in absence of ParB.

The transcription dynamics of *E. coli* RNAP was quantitatively assessed by a statistical analysis of transcription elongation and transcriptional pausing. Pause distributions were assessed using unbiased dwell time analysis^45^. The times needed for RNAP to transcribe through consecutive dwell time windows of five base pairs – defined as ‘dwell times’ – were calculated for all RNAP trajectories and used to construct a dwell time probability distribution function. The data was filtered to 1 Hz (moving average) for analysis of all data sets. The expected error (standard deviation) in the constructed distributions were estimated by bootstrapping the data 100 times^45^. To characterize the dwell time distributions, we divided it into three separate time ranges, as described in detail previously^45^: the pause-free transcription elongation region (0.1-1 s), which contained the elongation peak; the short elemental pause region (1-5 s); and the long backtrack pause region (>5 s). The elongation rate is given by 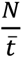, where *N* is the dwell time window size in bp, and 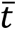 denotes the peak position of the gamma-fitted distribution. To calculate the probabilities of the short and long pauses, we integrated the dwell time distribution over the corresponding dwell time regions.

### Modelling and molecular dynamics simulations

We modelled the DNA as a semiflexible polymer made of 1,000 spherical beads of size *σ* = 5.5 *nm* = 16 *bp*. We placed *parS* in the middle of the chain where ParB beads are recruited and diffuse. The beads interact purely by excluded volume following the shifted and truncated Lennard-Jones (LJ) force field

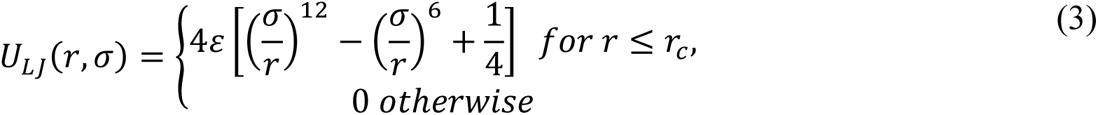

where *r* denotes the distance between any two beads and 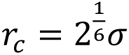 is the cut-off. We defined the bonds between two monomers along the DNA contour length by the finite extensible nonlinear elastic (FENE) potential, given by:

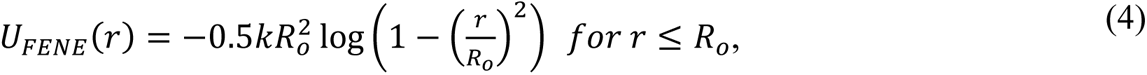

with *k* = 30 *ε*/*σ*^2^the spring constant and *R*_*o*_ = 1.5 *σ* the maximum length of the bond. We introduced the persistence length of the DNA chain as a bending potential energy between three consecutive beads given by:

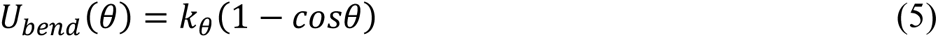

where *θ* is the angle between two bonds and *k*_*θ*_ = 10 *k*_*B*_*T* is the bending stiffness constant, corresponding to a persistence length of about 10 beads or ∼55 nm.

We performed the simulations in a constant-volume and constant-temperature (NVT) ensemble with implicit solvent utilizing the Langevin heat bath with local damping constant set to *γ* = 1 so that the inertial time equals the Lennard-Jones time 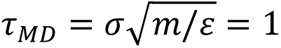 and *m* = 1 is the mass of each bead. We let equilibrate the chain for long enough to reach a constant radius of gyration, before turning on the ParB recruitment and diffusion. We evolved the Langevin equation in LAMMPS (*40*) using a Velocity-Verlet integration scheme and a timestep *dt* = 0.002.

We performed mapping to real time and length units by using the Brownian time 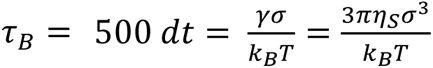. Using the viscosity of water *η_S_* = 1 *cP* we obtained that *τ_B_* = 0.5 *μs* We typically dump the positions of the beads every 10^4^*τ*_*B*_ = 5 *ms*. It’s also useful for later to note that with the coarse graining of 1 bead to 5.5 nm we also have that 1 *μm* = 182 *σ*.

We modelled ParB in the system by calling, within the LAMMPS engine, an external program that modifies the types of the beads to account for the loading and diffusion of ParB on DNA; in other words, ParB is implicitly modelled. At *t* = 0 we loaded a ParB protein onto the *parS* site and allowed it to diffuse with a constant 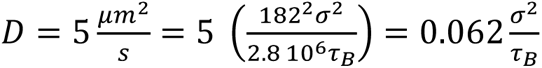 Note that this is 100x faster than the real diffusion rate of ParB, around 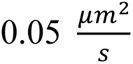. The diffusion is implemented by attempting the movement of a loaded ParB either to its left or to its right with probability 0.125 every *τ*_*B*_ = 500 *dt* timesteps (recall that MSD = 2 Dt in a 1D system, hence why the jump probability is twice the diffusion coefficient D). The diffusion cannot happen (the move is rejected) if the attempt brings a ParB protein either on top of another ParB or beyond the ends of the DNA.

On top of diffusion, we add a loading rate at ParS at a variable rate *κ*_*on*_ and an unloading process at fixed rate *κ*_*off*_ = 0.001 1/*τ*_*B*_, i.e. at a timescale *T*_*off*_ = 0.5 *ms*. This timescale is faster than the one seen experimentally but we argue that the overall behaviour of the system is controlled by the ratio of loading and unloading timescales. In line with our previous work on the recruitment mechanism of ParB proteins, we implement a stochastic recruitment at the same rate of the loading at ParS (*κ*_*on*_), with the difference that the recruitment probability is associated with each loaded ParB and can happen *in-cis* (with probability *p*_*c*_ = 0.083) or *in-trans* (with probability *p*_*T*_ = 1 − *p*_*C*_ = 0.9166), in such a way that the trans recruitment is 11 times larger than the cis recruitment. If the cis-recruitment is selected, one of the two 1D adjacent beads to a ParB is selected at random and, if unoccupied, a new ParB is added onto the chain. Otherwise, if the latter ‘*in-trans*’ mechanism is selected, we compute the list of 3D proximal neighbors which must be (i) within a Euclidean cutoff distance of 2*σ* = 11 *nm* and (ii) farther than the second-nearest neighbor in 1D (i.e. the first and second nearest neighbors cannot be picked). Once the list of 3D neighbors is compiled, we randomly pick one of these from the list (if not empty), load a new ParB protein, and resume the Langevin simulation.

### Implementation of Dimer-of-Dimers and Multimer-of-Dimers bridging

To precisely regulate the binding mechanism between ParB and control the valence of interactions we decorate each polymer bead with 2 patches at antipodal positions of the spherical bead. The central bead and the patches rotate as a rigid body and there is no constraint on the rotation of the patches, e.g. no dihedral potential is imposed. The interaction between patches is turned on only if the polymer bead is occupied by a ParB protein and we decide whether to turn one or both on. The former situation leads to the formation of one-to-one bridges (dimer-of-dimers) while the latter to the formation of multimers-of-dimers. The interaction between patches is regulated by a Morse potential of the form

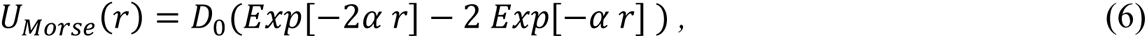

for *r* < *r*_*c*_ = 2 *σ* and 0 otherwise. *D*_0_is a parameter we vary in the simulations to explore the phase diagram, while *α* = 0.5 to ensure short-range interactions. We note that the cut off *r*_*c*_ = 2 *σ* = 11 *nm* is set on purpose to ensure that it is realistic for a pair-wise ParB interaction. The code to run this simulation can be found at https://git.ecdf.ed.ac.uk/taplab/parbcondensation.

## Acknowledgments

We thank Brian T. Analikwu for useful discussions.

## Author contributions

Conceptualization: MT, SG, CD; Fluorescence experiments: MT, TB; Formal analysis: MT, RB; Magnetic tweezers experiments and analysis: RJ; Atomic force microscopy experiments and analysis: AMG; Protein purification: HA; DNA constructs and cloning: MT, JvdT; Molecular dynamics simulations: DM; Visualization: MT, DM; Funding acquisition: DM, SG, CD; Supervision: DM, SG, CD; Writing – original draft: MT, CD; Writing – review & editing: all authors.

## Data and materials availability

All data are available in the main text or the supplementary material. The code used in molecular dynamics simulations is open-access and deposited at https://git.ecdf.ed.ac.uk/taplab/parbcondensation. All raw data from experiments are available upon request to corresponding authors.

## Delcaration of interests

Authors declare that they have no competing interests.

## Supplementary Materials

Figs. S1 to S11

**Table 1.** Plasmids and cloning strategy used in this study for creation of the DNA constructs.

|  |  |  |  |
| --- | --- | --- | --- |
| Plasmid Code | Plasmid Name |  |  |
| #179 | pBlueScript-GoldenGate-v2 |  |  |
| Cloning technique | Primers | Template | Template Origin |
| PCR (Q5 NEB E0555L) | JT492, JT 493 | #64 pBlueScript-GoldenGate | Cees Dekker lab |
| KLD reaction | NEB M0554S |  |  |
| Plasmid Code | Plasmid Name |  |  |
| #64 | pBlueScript-GoldenGate |  |  |
| Cloning technique | Primers | Template | Template Origin |
| PCR (Q5 NEB E0555L) | JT290, JT291 | #61 pBlueScript-noBsaI | Cees Dekker lab |
| KLD reaction | NEB M0554S |  |  |
| Plasmid Code | Plasmid Name |  |  |
| #61 | pBlueScript-noBsaI site |  |  |
| Cloning technique | Primers | Template | Template Origin |
| PCR (Q5 NEB E0555L) | JT288, JT298 | #18 pBlueScriptII SK+ | Stratagene |
| KLD reaction | NEB M0554S |  |  |
| Plasmid Code | Plasmid Name |  |  |
| #66 | pGGA-JT292JT293 |  |  |
| Cloning technique | Primers | Template | Template Origin |
| PCR (Q5 NEB E0555L) | JT492, JT493 | Lambda | NEB N3011S |
| PCR (Q5 NEB E0555L) | JT403, JT401 | pGGAselect | NEB N0309AAVIAL |
| Blunt ligation | Enzymes used: T4 PNK (NEB M0201), DPNI (NEB R0176), T4 DNA Ligase (NEB M0202) |  |  |
| Plasmid Code | Plasmid Name |  |  |
| #67 | pGGA-JT294JT295 |  |  |
| Cloning technique | Primers | Template | Template Origin |
| PCR (Q5 NEB E0555L) | JT294, JT295 | Template GeneBlock34 | IDT Technologies |
| PCR (Q5 NEB E0555L) | JT403, JT401 | pGGAselect | NEB N0309AAVIAL |
| Blunt ligation | Enzymes used: T4 PNK (NEB M0201), DPNI (NEB R0176), T4 DNA Ligase (NEB M0202) |  |  |
| Plasmid Code | Plasmid Name |  |  |
| #69 | pGGA-JT310JT314JT311JT315 |  |  |
| Cloning technique | Primers | Template | Template Origin |
| PCR (Q5 NEB E0555L) | JT310, JT314 | Lambda | NEB N3011S |
| PCR (Q5 NEB E0555L) | JT315, JT311 | Lambda | NEB N3011S |
| PCR (Q5 NEB E0555L) | JT403, JT401 | pGGAselect | NEB N0309AAVIAL |
| Gibson Assembly | NEB E5520 (NEB HiFi mix) |  |  |
| Plasmid Code | Plasmid Name |  |  |
| #163 | pGGA-T7A1-ParS |  |  |
| Cloning technique | Primers | Template | Template Origin |
| PCR (Q5 NEB E0555L) | JT472, JT473 | #65 pGGA-JT298JT299 | NEB N3011S |
| PCR (Q5 NEB E0555L) | JT466, JT467 | #126 pSC-T7A1reverse+parS | NEB N3011S |
| Gibson Assembly | NEB E5529 (NEB HiFi mix) |  |  |
| Plasmid Code | Plasmid Name |  |  |
| #65 | pGGA-JT298JT299 |  |  |
| Cloning technique | Primers | Template | Template Origin |
| PCR (Q5 NEB E0555L) | JT298, JT299 | Template GeneBlock35 | IDT Technologies |
| PCR (Q5 NEB E0555L) | JT403, JT401 | pGGAselect | NEB N0309AAVIAL |
| Blunt ligation | Enzymes used: T4 PNK (NEB M0201), DPNI (NEB R0176), T4 DNA Ligase (NEB M0202) |  |  |
| Plasmid Code | Plasmid Name |  |  |
| #71 | pGGA-JT312JT316JT313JT317 |  |  |
| Cloning technique | Primers | Template | Template Origin |
| PCR (Q5 NEB E0555L) | JT312, JT316 | Lambda | NEB N3011S |
| PCR (Q5 NEB E0555L) | JT313, JT317 | Lambda | NEB N3011S |
| PCR (Q5 NEB E0555L) | JT403, JT401 | pGGAselect | NEB N0309AAVIAL |
| Gibson Assembly | NEB E5520 (NEB HiFi mix) |  |  |
| Plasmid Code | Plasmid Name |  |  |
| #126 | pSuperCos-T7A1reverse+parS |  |  |
| Cloning technique | Primers | Template | Template Origin |
| PCR (Q5 NEB E0555L) | MT236, MT239 | #92 pSC-T7A1reverse | NEB N3011S |
| PCR (Q5 NEB E0555L) | MT235, MT238 | #92 pSC-T7A1reverse | NEB N3011S |
| PCR (Q5 NEB E0555L) | MT234, MT237 | #92 pSC-T7A1reverse | NEB N3011S |
| Gibson Assembly | NEB E5520 (NEB HiFi mix) |  |  |
| Plasmid Code | Plasmid Name |  |  |
| #92 | pSC-T7A1reverse |  |  |
| Cloning technique | Primers | Template | Template Origin |
| PCR (Q5 NEB E0555L) | JT405, JT406 | #73 pIA146 T7A1 | Irina Artsimovitch lab |
| Digest | #29 pSuperCos1-Lambda1,2 | MscI | NEB R0534S |
| Blunt Ligation | Enzymes used: T4 PNK (NEB M0201), DPNI (NEB R0176), T4 DNA Ligase (NEB M0202) |  |  |
| Plasmid Code | Plasmid Name |  |  |
| #29 (from Ref. <sup>70</sup>) | pSuperCos1-Lambda1,2 |  |  |
| Cloning technique | Primers | Template | Template Origin |
| Step 1 |  |  |  |
| PCR (Q5 NEB E0555L) | 56-9,5kblambdafw, 57-9,5kblambdarev | Lambda | NEB N3011S |
| Digest | Digest PCR product with NotI + PspOMI, ligate in pSuperCos1 (Agilant) digested with NotI |  |  |
| Step 2 |  |  |  |
| PCR (Q5 NEB E0555L) | 58-5kblambdafw, 59-5kblambdarev | Lambda | NEB N3011S |
| Digest | Digest PCR product and plasmid made in step2 with BglII + BssHII |  |  |
| Ligation | T4 DNA Ligase (NEB M0202) |  |  |
| Plasmid Code | Plasmid Name |  |  |
| #186 |  |  |  |
| Cloning technique | Primers | Template | Template Origin |
| PCR (Q5 NEB E0555L) | JT494, JT495 | #18 pBlueScriptII SK+ | Stratagene |
| KLD reaction | NEB M0554S |  |  |

**Table 2.**
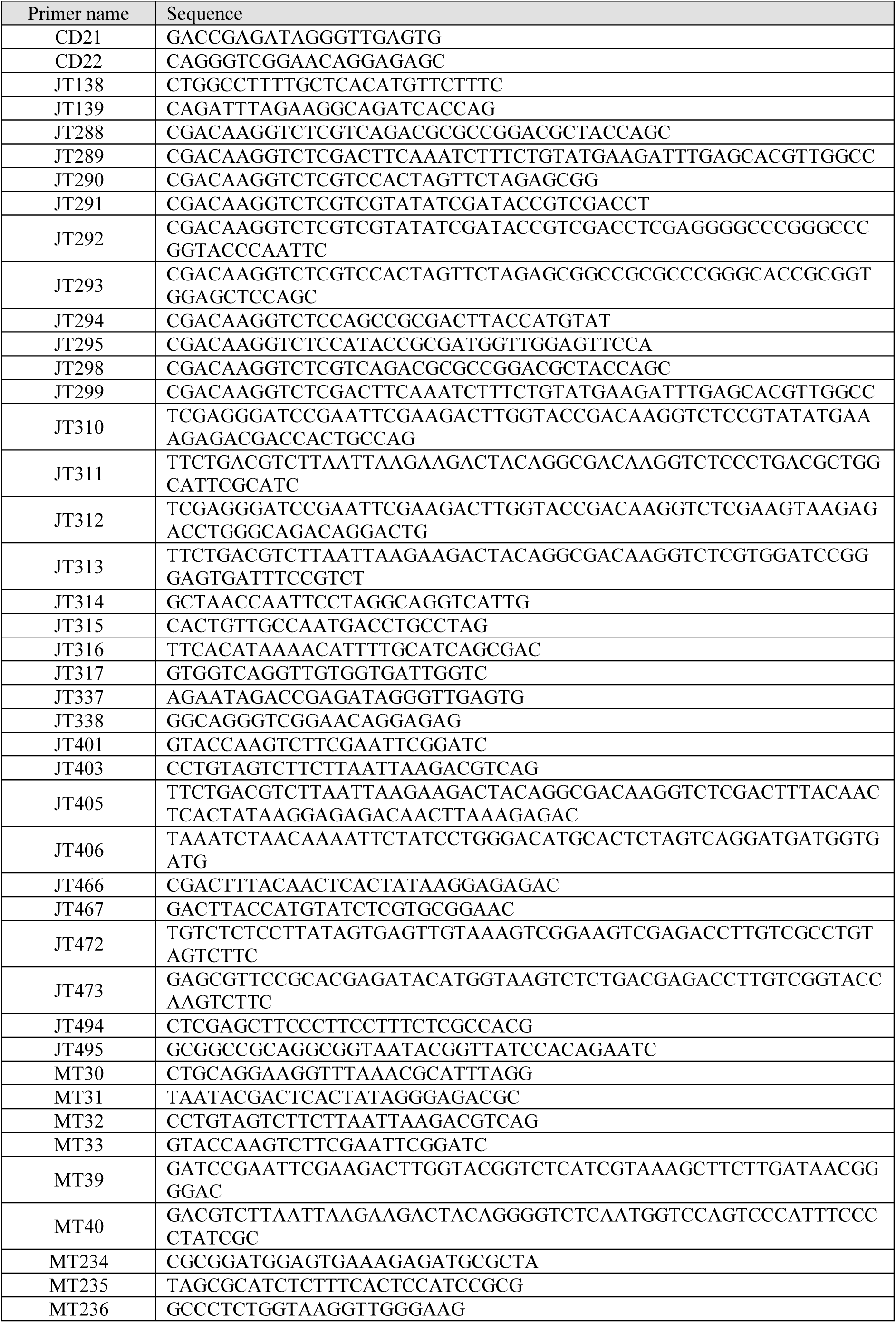
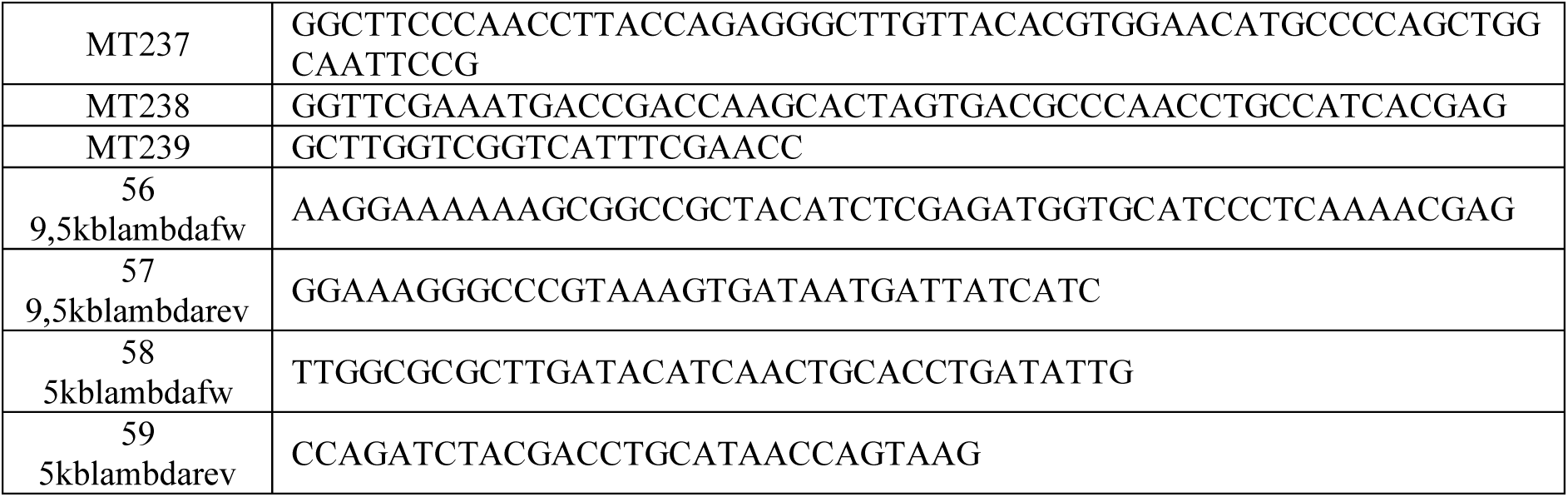
DNA primers used in this study for cloning and plasmid modification.

**Table 3.**
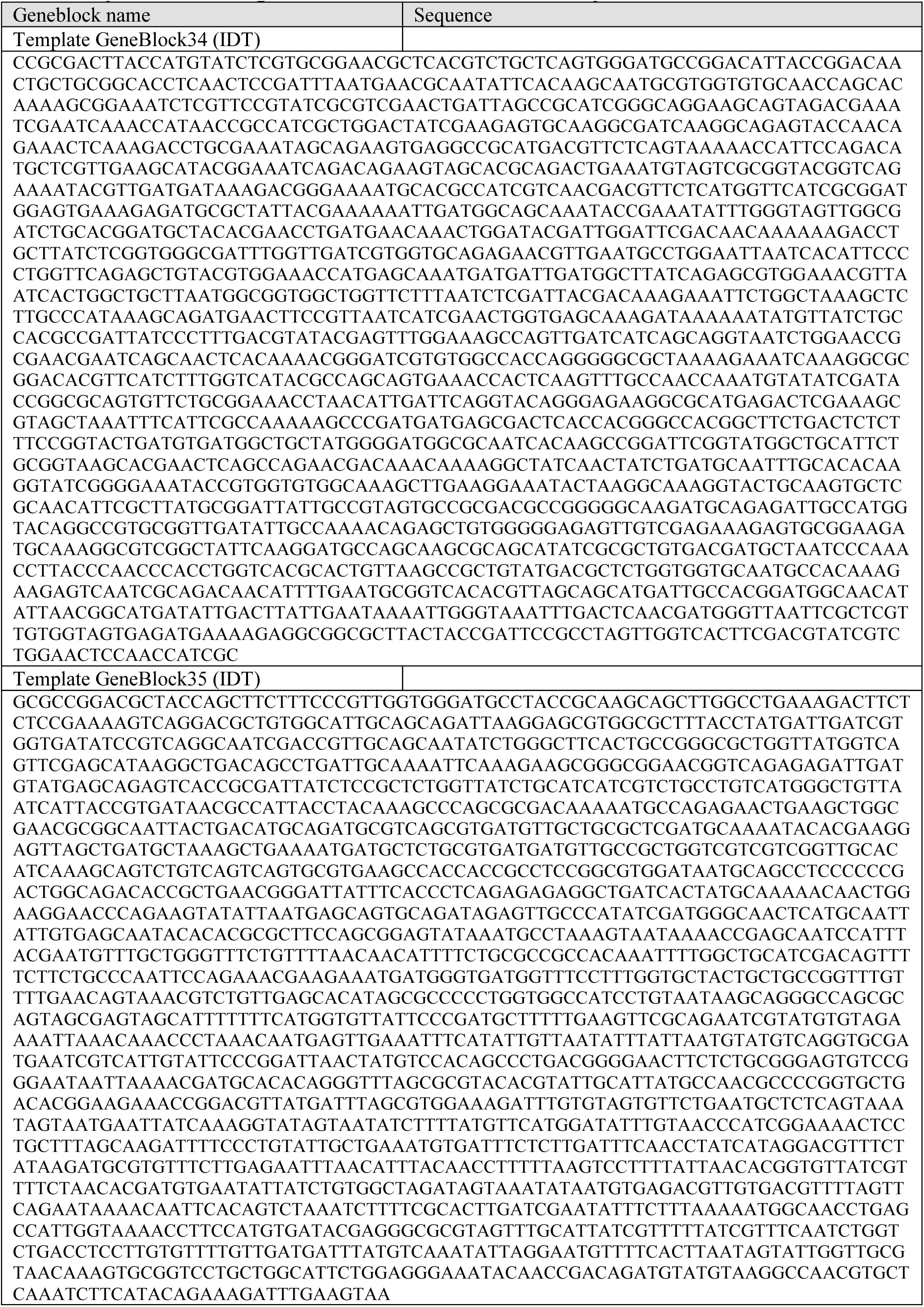
Synthetic DNA gene constructs used in this study.

## Supplementary movies

**Movie S1. High-speed AFM movie of ParB loading and transiently looping the DNA molecule.**

**Movie S2. Molecular dynamics simulations of DNA polymer conformation in presence of ParB (*ϵ* = 5 and *κ* = 20) with dimer-of-dimers interaction (DoD).**

**Movie S3. Molecular dynamics simulations of DNA polymer conformation in presence of ParB (*ϵ* = 16 and *κ* = 20) with dimer-of-dimers interaction (DoD).**

**Movie S4. Molecular dynamics simulations of DNA polymer conformation in presence of ParB (*ϵ* = 10 and *κ* = 10) with multimer-of-dimers interaction (MoD).** The polymer collapses in the condensed state maintained via transient ParB-ParB bridges.

**Movie S5. Molecular dynamics simulations of DNA polymer conformation in presence of ParB (*ϵ* = 2 and *κ* = 20) with multimer-of-dimers interaction (MoD).** The polymer remains in the extended state even when ParB proteins are spread over the entire length of the polymer.

