## Supplementary figures for "Dynamic ParB-DNA interactions initiate and maintain a partition condensate for bacterial chromosome segregation"

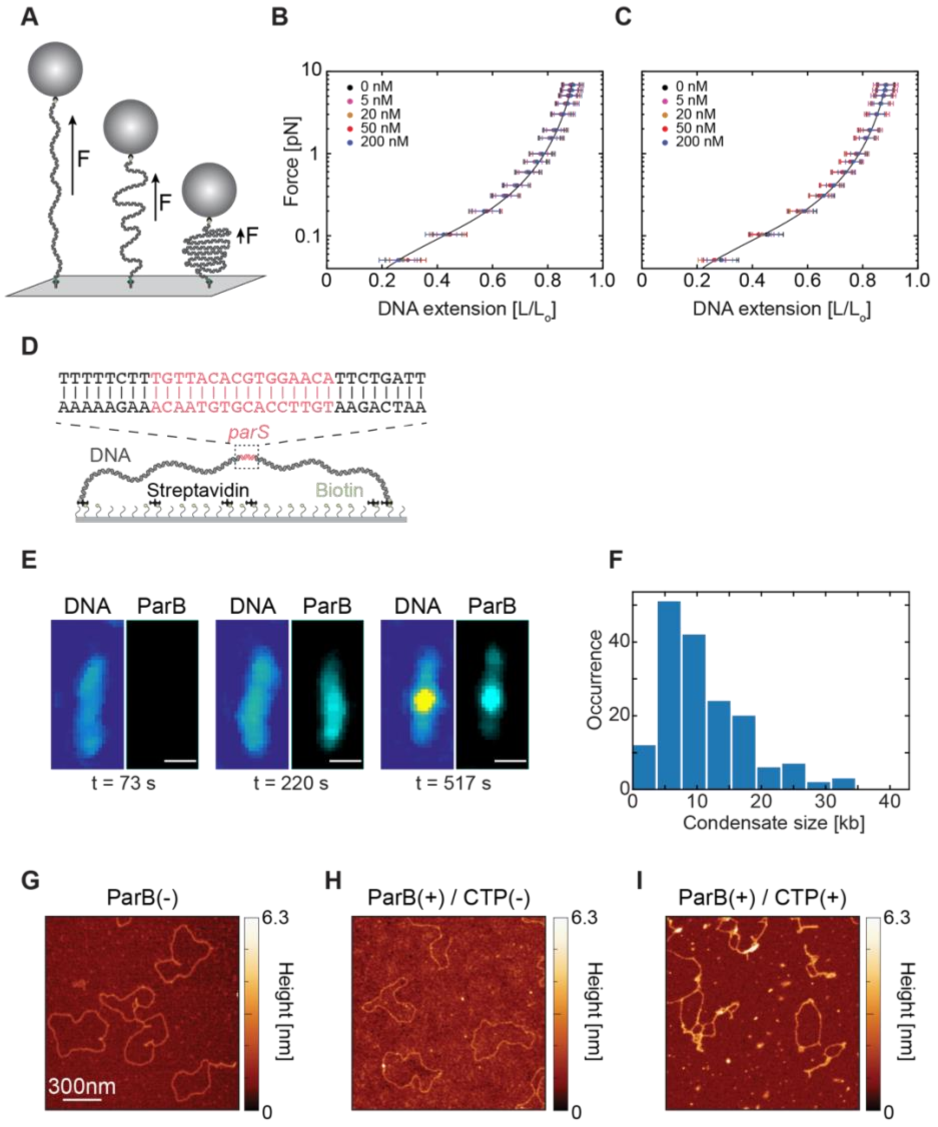


**Figure S1. ParB proteins do not condense the DNA in the absence of CTP or without a *parS* site.**

**(A)** Representation of experimental MT assay where the force is gradually lowered and the bead tethered to 12 kbp linear DNA moves towards the surface in presence of different ParB concentrations. **(B)** Average DNA extension (mean±SD) at lowering forces (N; 0 nM: 61, 5 nM: 61, 20 nM: 60, 50 nM: 60, 200 nM: 57) in the absence of a *parS* site on the DNA. (**C)** Average DNA extension (mean±SD) at lowering forces (N; 0 nM: 16, 5 nM: 15, 20 nM: 15, 50 nM: 17, 200 nM: 11) in the absence of CTP nucleotide. Black lines in (B) and (C) represent a WLC fit to the 0 nM ParB data. **(D)** Schematic representation of $DNA_{parS}$ tethered with both ends to a glass surface via streptavidin-biotin interactions (see Methods). (**E)** Representative fluorescence images of $DNA_{parS}$ and $ParB^{TMR}$ signals at three time points during the experiment: (left) before ParB is present, (middle) at the moment that ParB covers the $DNA_{parS}$, and (right) after the $DNA_{parS}$ is visibly condensed by ParB molecules. (**F)** The size of ParB-induced DNA condensates on the 42 kbp $DNA_{parS}$ molecule (N = 161). (**G)** Dry AFM image of 4.2 kbp circular $DNA_{parS}$ molecule in the absence of ParB proteins. (**H)** Same as in (D) but in the presence of 5 nM ParB proteins. (**I)** Same as (D) but in the presence of both 1 mM CTP and 5 nM ParB. Related to Figure 1.

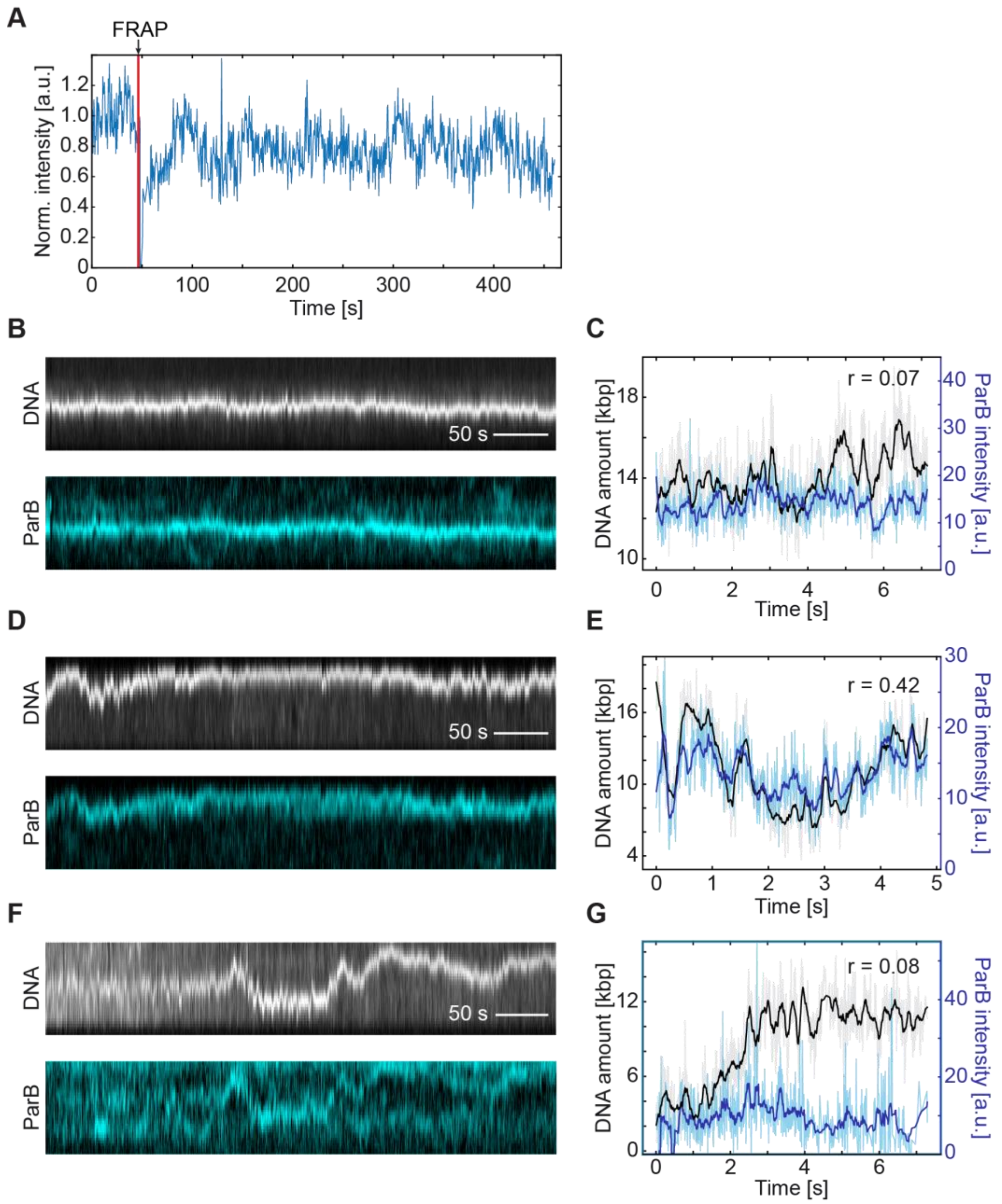


**Figure S2. ParB:DNA clusters are dynamic and experience high ParB turnover. (A)** Example FRAP (fluorescence recovery after photobleaching) fluorescence intensity plot of a single DNA$_{parS}$ molecule from Fig. 2A. The bleaching event is marked with red line. (**B)** Kymographs of DNA$_{parS}$ (top, grey) and ParB$^{alexa647}$ (bottom, cyan) after the bleaching recovery events. (**C)** Quantification of the ParB signal intensity (light blue: raw data; purple: filtered data (Savitzky-Golay, time window =11 frames)) and the amount of DNA within the condensed region of the DNA$_{parS}$ (light gray: raw data; black: filtered data). Pearson correlation coefficient (r) shown in legend. (**D-E)** Same as (B) and (C) for an example molecule with a higher r value. (**F-G)** Same as (B) and (C) for an example molecule with a lower r value. Related to Figure 2.

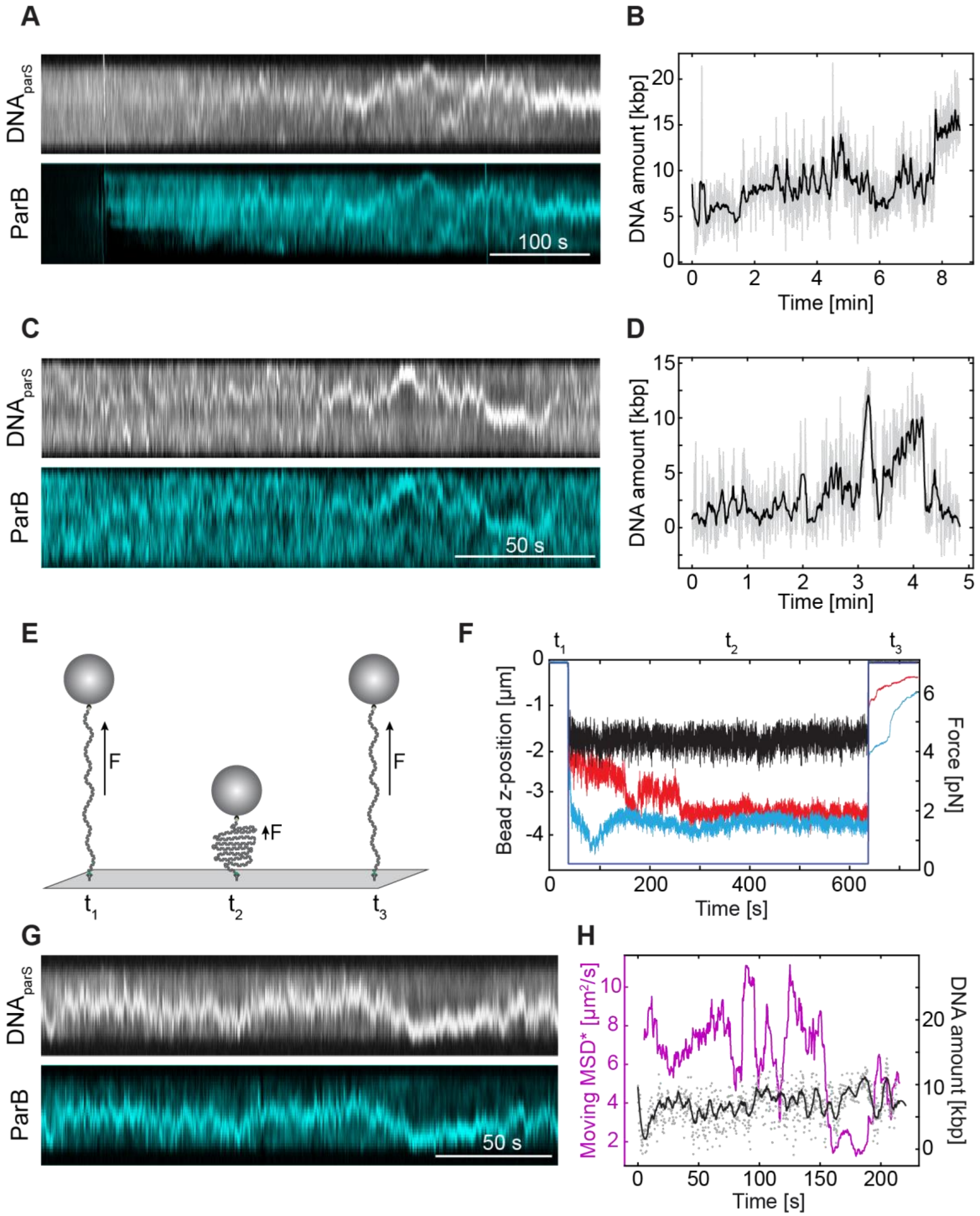


**Figure S3. DNA$_{parS}$ molecules experience highly dynamic and fluctuating DNA condensation in the initial stages. (A)** Time course experiment of ParB inflow and ParB-induced DNA condensation. Kymographs of DNA$_{parS}$ (top, grey) and ParB$^{alexa647}$ (bottom, cyan) from the beginning of ParB loading up to dynamic DNA condensation. (**B)** Quantification of the amount of DNA within the condensed region of the $DNA_{parS}$ (light grey: raw data; black: filtered data (Savitzky-Golay, time window =11 frames)). **C-D)** Same as in (A) and (B). (**E)** Time course experiment in MT assay. The DNA bead is lowered from 5 pN ($t_1$) to 0.3 pN for 5 min ($t_2$), and then back to 5 pN ($t_3$). (**F)** Magnetic bead z-position representing $DNA_{parS}$ extension in the presence of 25 nM ParB. Black trace without ParB does not show DNA condensation events, while in presence of ParB the $DNA_{parS}$ experiences diverse discrete condensation steps (blue, red). (**G)** Kymographs of $DNA_{parS}$ (top, grey) and $ParB^{alexa647}$ (bottom, cyan) after ~20 min post ParB inflow, during the stable condensation stage. (**H)** Fluctuating DNA cluster position during a stable condensation. Magenta - apparent Mean Squared Displacement (MSD*, window size = 52 frames) of the ParB:DNA condensate tracked by the relative position along the 1D line of the tethered DNA. Gray dots – raw data of the DNA amount within the ParB:DNA condensate in each frame. Black – Savitzky-Golay filter (window size = 11 frames) of the DNA amount withing the condensate in shown in grey. Related to Figure 2.

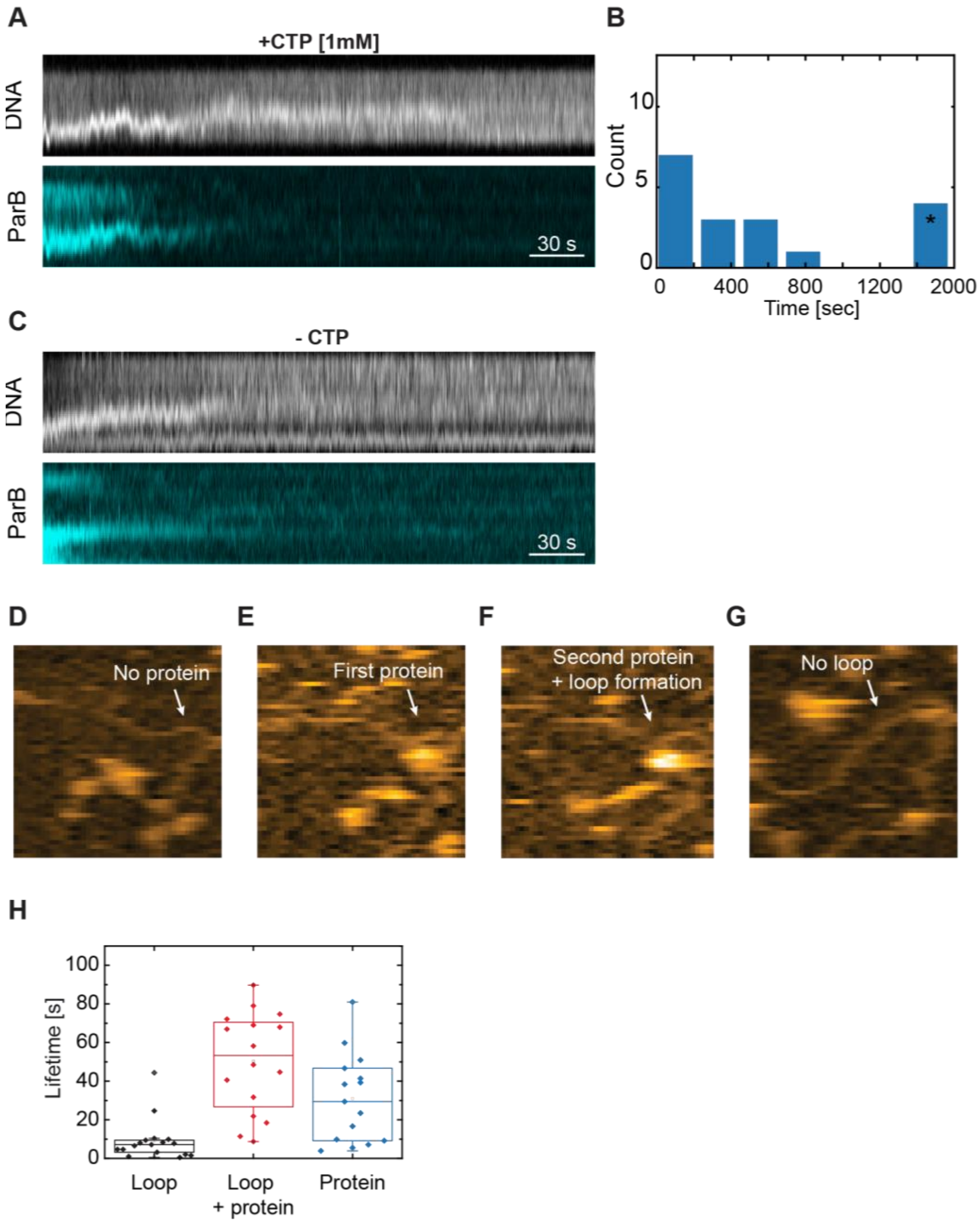


**Figure S4. ParB-induced DNA condensation is reversible in the presence and absence of CTP.**

**(A)** Kymographs of DNA$_{parS}$ (top, grey) and ParB$^{TMR}$ (bottom, cyan) from the moment of buffer wash that does not contain ParB$^{TMR}$ molecules. (**B**) Time from the beginning of the washing until the condensate entirely disassembles. Starred bin represents condensates that did not wash during the entirety of the video. (**C**) Same as (A) for the washing experiment using the buffer containing neither CTP nor ParB molecules. **(D-g)** High-speed AFM snapshots of transient DNA loop formation in the presence of ParB (7 nM; see Movie S1). (**H**) Loop lifetimes on the DNA$_{parS}$ in the absence (black; mean±SD: 5.6±3.3 s; N = 15) and presence (red; mean±SD: 50±25 s; N = 16) of ParB molecules, and protein lifetime under the same conditions (blue; mean±SD: 30.8±22.9 s; N = 15). Related to Figure 3.

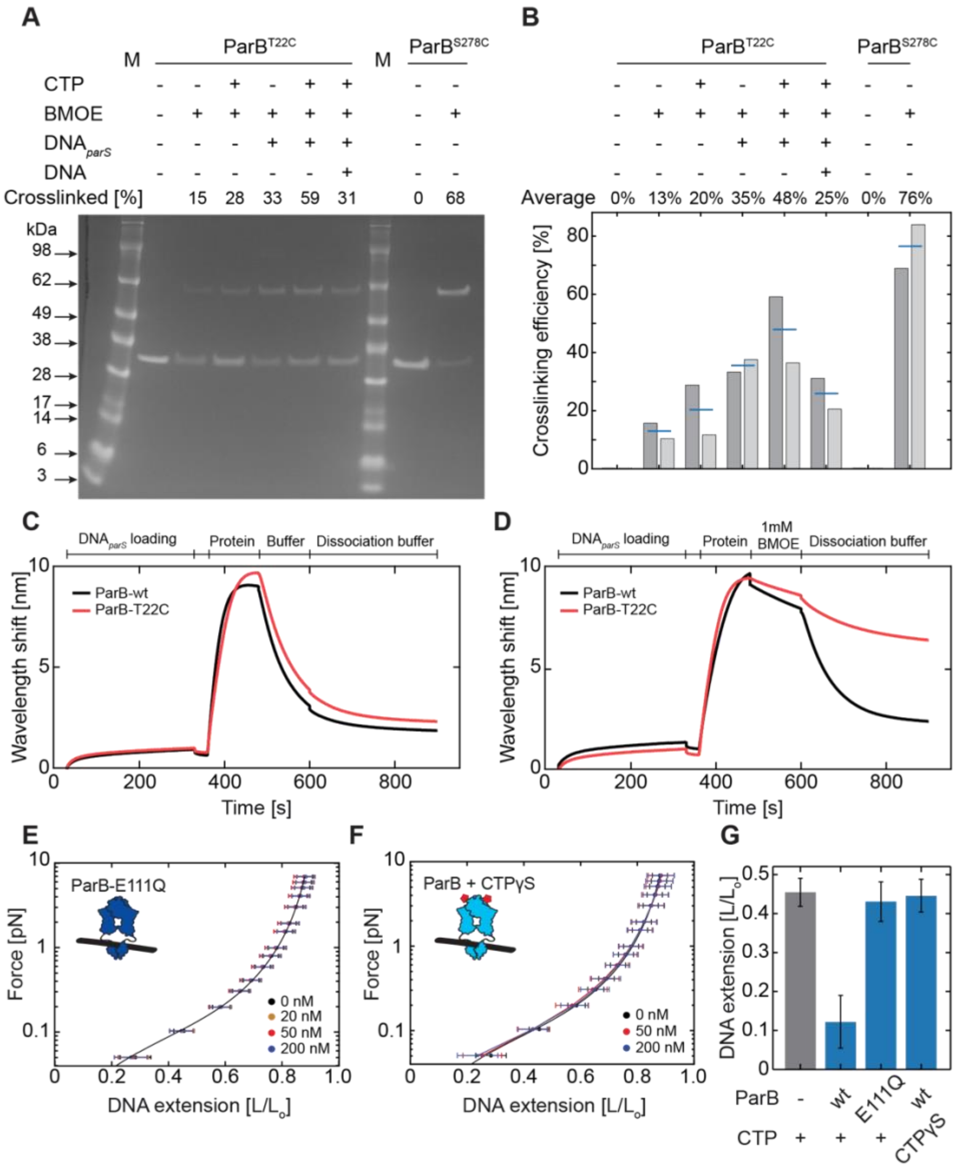


**Figure S5. ParB mutants (T22C and S278C) can be crosslinked at their C-termini by BMOE.**

**(A)** SDS-PAGE gel showing crosslinking efficiency of ParB$^{\mathbf{T22C}}$ (lanes 2-7) and ParB$^{\mathbf{S278C}}$ lanes (9-10) in the presence of 1 mM BMOE. Crosslinking efficiency is shown in the percentage of the total signal from both bands. Experimental conditions shown on top and more detailed in the Methods section. (**B**) Crosslinking efficiency from a duplicate experiment under the same conditions. Blue lines represent the average crosslinked fraction from two experiments. (**C**) DNA loading and dissociation of ParB-wt and ParB$^{\mathbf{T22C}}$ (1 μM) on DNA$_{parS}$ measured by BLI in the presence of 1 mM CTP (see Methods). During dissociation, an equivalent buffer lacking ParB protein and CTP was applied. (**D**) Same as in (C), with the additional step of introducing 1 mM BMOE for cysteine-crosslinking between the loading and dissociation step. (**E**) Average DNA$_{\mathrm{parS}}$ extension (mean±SD) in MT in the presence of catalytically inactive ParB$^{\mathbf{E111Q}}$ mutant (N; 0 nM: 32, 20 nM: 36, 50 nM: 39, 200 nM: 36). **F)** Same as in (E), but in the presence of the wild-type ParB protein with 1 mM CTPγS (N; 0 nM: 32, 50 nM: 33, 200 nM: 32). **(G)** Relative DNA extension in different conditions at an applied force of F = 0.1 pN. Related to Figure 4.

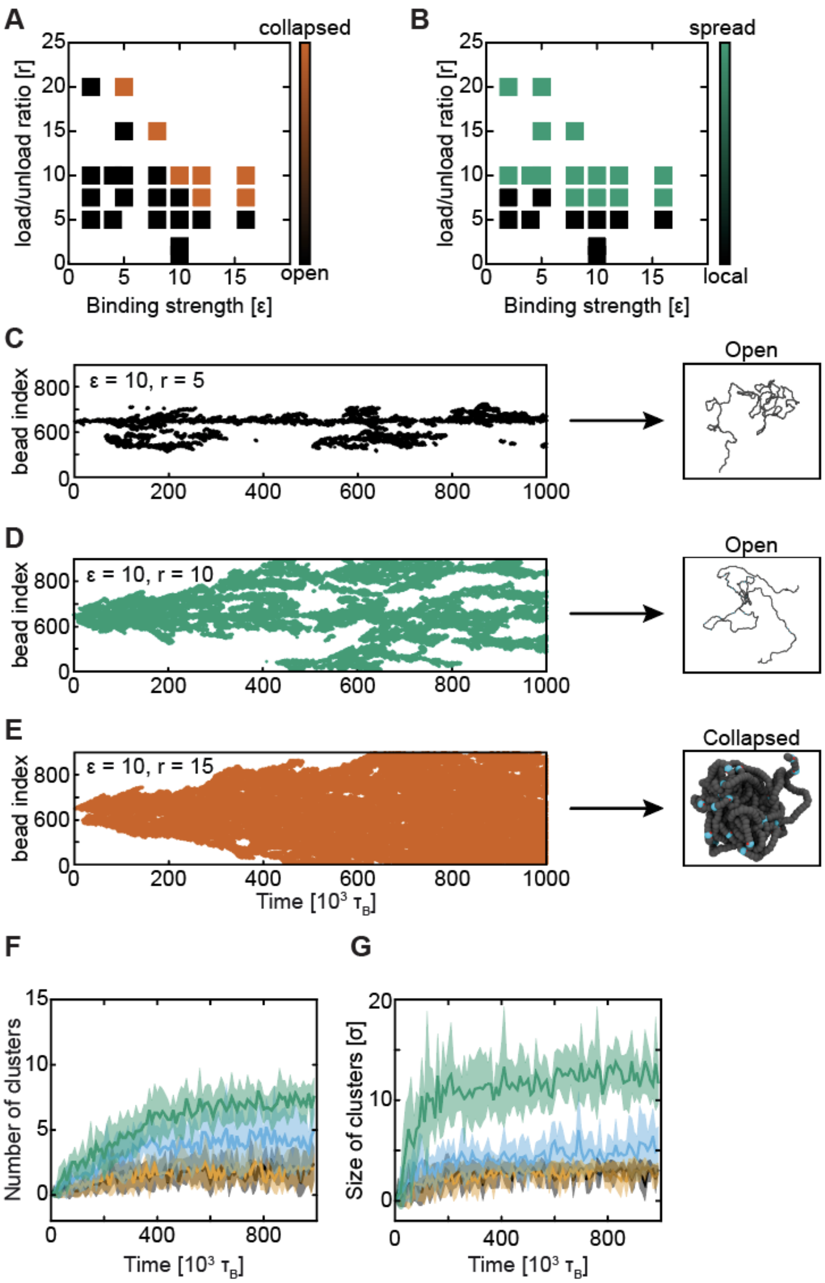


**Figure S6. Efficient DNA condensation requires spreading and sufficient ParB interaction strength**

**(A)** Final DNA polymer conformation (collapsed or open) in relation to load/unload rate (κ) and binding strength (epsilon). **(B)** ParB distribution over the DNA polymer (local or spread) in relation to load/unload rate (k) and binding strength (ε). **(C)** Left - Kymograph showing ParB distribution along the simulated DNA polymer at ε = 10 and κ = 5. Right – a snapshot of the final polymer conformation. **(D-E)** Same as (C) for k = 10 and κ = 15, respectively. **(F)** The number of DNA clusters over time per DNA. Legend shown on top. κ = 10. Shaded area is SD among 10 replicas of the same system with different initial configuration and seed. **(G)** The size of individual DNA clusters over time within a single DNA. Legend same as in F). κ = 10. Related to Figure 5.

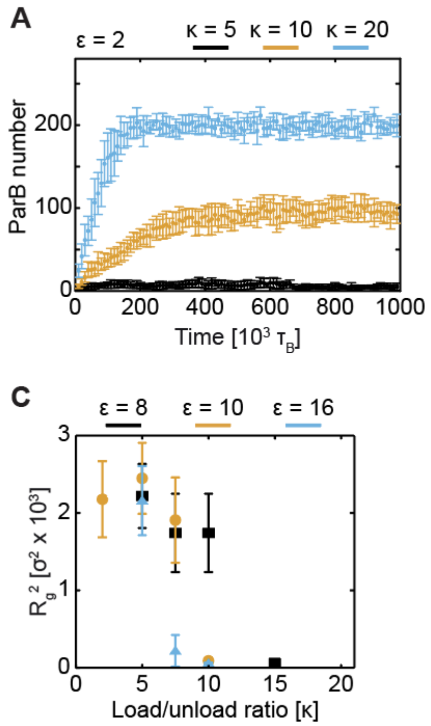


**Figure S7. DNA-condensation by ParB experiences switch-like behavior due to a high ParB cooperativity.**

**(A)** Total ParB numbers over time at increasing load/unload rates (κ) at a fixed interaction strength of ε = 2. (**B)** Same as (A) for ε = 10. **(C)** Radius of gyration of simulated DNA polymers as a function of κ. Related to Figure 5.

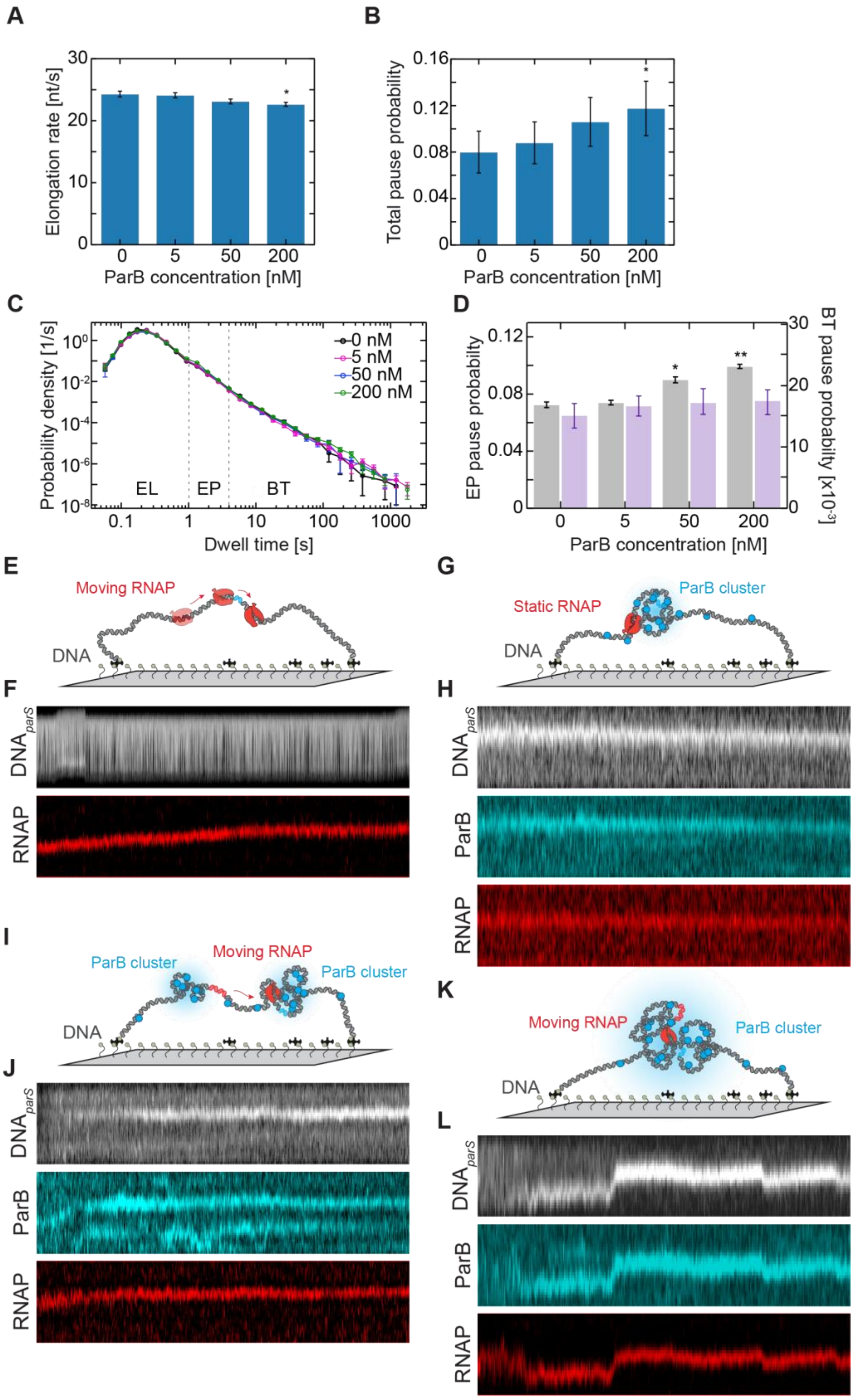

A
B
Elongation rate [nt/s]
ParB concentration [nM]
Total pause probability
C
D
Probability density [1/s]
Dwell time [s]
0 nM
5 nM
50 nM
200 nM
EL
EP
BT
EP pause probability
BT pause probability [x10-3]
E
G
Moving RNAP
DNA
Static RNAP
ParB cluster
F
H
DNAparS
RNAP
ParB
I
K
J
L

**Figure S8. ParB proteins can efficiently condense the DNA in the presence of transcribing RNAP**

**(A)** Pause-free elongation rate (mean±SD) and (**B**) total transcription pause probability (mean±SD) in the presence of increasing concentrations of ParB, extracted from Dwell time analysis in (C). (**C)** Superimposed dwell time distributions from RNAP trajectories in the presence of increasing concentrations of ParB (N; 0 nM: 27, 5 nM: 46, 50 nM: 29, 200 nM: 51). The distribution was constructed using the dwell times needed for RNAP to transcribe successive 5-bp segments of DNA. The dwell time distributions are separated with boundaries (see also Refs. 1,2) for the elongation region (0.1-1 s), short elemental pauses EP (1-5 s), and backtrack-induced pauses BT (>5 s). (**D**) The probability of elemental pause EP (grey) increases significantly with ParB concentration, while long backtrack pauses (purple) are not affected by ParB. Data was subject to statistical analysis using an unpaired, two-tailed t-test (p: ** <0.01; *< 0.05). **(E)** Schematic representation of RNAP transcription in our fluorescence assays **(F)** Kymographs of $DNA_{parS/T7A1}$ (top, grey) and transcribing $RNAP^{Alexa647}$ (bottom, red) from the moment of transcription initiation (**G)** Schematic representation of ParB:$DNA_{parS/T7A1}$ cluster in presence of non-transcribing RNAP (**H)** Kymographs of $DNA_{parS/T7A1}$ (top, grey), $ParB^{TMR}$ (bottom, cyan) and non-transcribing/halted $RNAP^{Alexa647}$ (bottom, red). (**I)** Schematic representation of ParB:$DNA_{parS/T7A1}$ cluster in presence of transcribing RNAP at higher end-to-end length of $DNA_{parS/T7A1}$. (**J)** Same as (H). (**K)** Schematic representation of ParB:$DNA_{parS/T7A1}$ cluster in presence of transcribing RNAP at lower end-to-end length of $DNA_{parS/T7A1}$. (**L)** Same as (H). Related to Figure 6.
